# Species-specific iNOS expression distinguishes epithelial and myeloid IFNγ responses in tuberculosis

**DOI:** 10.64898/2026.01.11.697682

**Authors:** Fabian Stei, Erin F. McCaffrey, Björn Zessin, Jonathan Pioch, Stefan Simm, Beate Stubbe, Ralf Ewert, Frank Verreck, Bianca E Schneider, Anca Dorhoi, Bryan D Bryson, Björn Corleis

**Affiliations:** Institute of Immunology, Friedrich-Loeffler-Institut, Greifswald - Isle of Riems, Germany; Spatial Immunology Unit, T-Lymphocyte Biology Section, Laboratory of Parasitic Diseases, National Institute of Allergy and Infectious Diseases, National Institutes of Health, Bethesda, MD, USA; Institute of Bioanalytics, University of Applied Sciences Coburg, Germany; Internal Medicine B, Pneumology, University Hospital Greifswald, Greifswald, Germany; Biomedical Primate Research Centre, Rijswijk, The Netherlands; Priority Area Infections, Research Center Borstel-Leibniz Lung Center, Germany; Ragon Institute of MGH, Harvard, and MIT, Cambridge, USA

## Abstract

**Rationale:** Tuberculosis vaccines aim to elicit protective immunity, with IFNγ-producing CD4⁺ T cells considered central mediators of host defense. In murine models, IFNγ activates macrophages to induce inducible nitric oxide synthase (iNOS), resulting in nitric oxide–dependent control of *Mycobacterium tuberculosis* (*Mtb*).

**Objective:** To define the relevance of IFNγ-induced iNOS activity in humans and other natural host of virulent mycobacterial species.

**Methods:** We systematically compared IFNγ-induced effector responses in *Mtb*-infected myeloid cells across species. Using bulk RNA sequencing, functional infection assays, and nitric oxide measurements, we assessed IFNγ responsiveness in human and mouse macrophages. These analyses were extended to monocytes from seven mammalian species and complemented by reanalysis of publicly available single-cell RNA-sequencing datasets and spatial proteomic imaging of tuberculous granulomas.

**Measurements and Main Results:** Single-cell transcriptomic reanalysis revealed minimal *NOS2* expression in myeloid cells from human and non-human primate granulomas. *In vitro*, IFNγ pretreatment failed to induce *NOS2* transcription, iNOS activity, or *Mtb* growth restriction in human macrophages, in stark contrast to murine cells. Across species, iNOS activity was largely restricted to mice, with limited induction in cattle monocytes. Instead, human respiratory epithelial cells consistently expressed *NOS2*, and multiplexed ion beam imaging localized iNOS protein to epithelial compartments adjacent to granulomatous lesions.

**Conclusion:** IFNγ signaling is uncoupled from iNOS induction in primate myeloid cells and epithelial compartments represent dominant sources of iNOS in human tuberculosis. These findings, challenge murine macrophage–centric paradigms and IFNγ-based correlates used in TB vaccine development and central to pathophysiology of tuberculosis and other pneumonias.

## Introduction

Tuberculosis (TB) remains the leading cause of death from a single infectious agent worldwide. The development of new vaccines is hindered by the lack of a robust correlate of protection (1). It is generally accepted that cellular immune responses—especially CD4⁺ IFNγ producing T helper 1 (Th1) cells—are beneficial in both humans and animal models (2). In humans, the importance of IFNγ is supported by studies of individuals with genetic deficiencies (Mendelian susceptibility to mycobacterial disease) in IFNγ or components of Th1 signaling pathways, who exhibit heightened susceptibility to mycobacteria (3–6). Similarly, the AIDS co-pandemic has underscored the relevance of Th1 immunity, as HIV-1-mediated depletion of CD4⁺ T cells is associated with increased TB susceptibility (7–9). In mouse models of TB, genetic deletion of IFNγ or Th1 cells results in profound susceptibility to infection with virulent *Mycobacterium tuberculosis* (*Mtb*) strains (10–13). Mouse models have also been instrumental in dissecting the mechanisms of IFNγ mediated TB control. Macrophages, the primary host cells for *Mtb*, express the IFNγ receptor (IFNGR), and IFNγ signaling induces the expression of antimicrobial effectors (14, 15). A key effector is iNOS, whose expression in macrophages leads to nitric oxide production within *Mtb*-containing phagosomes. Nitric oxide is thought to contribute directly to bacterial killing and to modulate downstream immune responses through S-nitrosylation of host proteins (16–24). In contrast to the well-defined role of IFNγ and iNOS in mice, the contribution of this pathway in human macrophages is less clear. Several studies have failed to detect *Mtb* growth inhibition in IFNγ stimulated human macrophages (25–30), unless cells were cultured under specialized conditions (31, 32). *In vivo,* iNOS protein has been detected at the outer rim of granuloma structures, and nitric oxide has been measured in the exhaled air of TB patients (33–36). However, most *in vitro* studies fail to detect NOS2 transcription or iNOS activity in IFNγ-stimulated human macrophages (37–40). Thus, while IFNγ is clearly relevant for human TB immunity, its direct antimicrobial impact on *Mtb*-infected human macrophages remains unresolved.

Recent advances in single-cell transcriptomics and spatial tissue profiling now allow direct interrogation of effector programs across species and cellular compartments within TB granulomas. Here, we combined controlled *in vitro* infection models with reanalysis of publicly available single-cell RNA-sequencing datasets to systematically examine IFNγ-induced iNOS expression across different species permissive to TB. We show that myeloid cells from humans, non-human primates, and other mammalian species lack *NOS2* transcription and functional iNOS activity despite strong IFNγ-driven inflammatory responses, in stark contrast to murine macrophages. Instead, iNOS expression localizes predominantly to epithelial compartments in both human and primate respiratory tissues, including in proximity to granulomatous lesions. These findings demonstrate a conserved uncoupling of IFNγ signaling from iNOS induction in primate myeloid cells and challenge the assumption that murine nitric oxide–dependent macrophage mechanisms are universally operative in TB immunity. Our results underscore the need to re-evaluate IFNγ-based correlates and extrapolation from experimental models in rational TB vaccine development.

## Methods

### Ethical approval

All biological sample collections were conducted under appropriate ethical oversight and in accordance with institutional and national guidelines. Human peripheral blood mononuclear cells (PBMCs) were isolated from anonymous buffy coats obtained through the University Hospital Greifswald blood donation center, with written informed consent and approval by the local ethics committee (BB 014/14). Bronchoalveolar lavage (BAL) fluid and airway biopsy samples were obtained as excess clinical material with informed consent (BB 058/19). Livestock blood samples were collected at the Friedrich-Loeffler-Institut under approved protocols (LALLF 7221.3-2-041/17; 7221.3-2-001/23). Mouse tissues were obtained post-mortem in accordance with German Animal Welfare regulations (§4 Absatz 3). Non-human primate PBMCs were obtained from the Biomedical Primate Research Centre under institutional approval (DEC). Detailed ethical information is provided in the Supplementary Methods.

### Study design

This exploratory study examined interspecies differences in IFNγ-mediated macrophage responses to *Mtb*. Human monocyte-derived macrophages and mouse bone marrow–derived macrophages were analyzed in parallel using bulk RNA sequencing and functional assays. Additional species (cattle, swine, goat, sheep, and non-human primates) were included for comparative analyses. Experiments were not blinded, and sample sizes were determined by availability. Replicate numbers and statistical tests are reported in the figure legends.

### Cell isolation and macrophage differentiation

PBMCs were isolated by density gradient centrifugation and used for monocyte enrichment and macrophage differentiation. Human macrophages were differentiated from CD14⁺ monocytes, and mouse macrophages were generated from bone marrow precursors. Cells were maintained under standardized culture conditions and stimulated with species-specific recombinant IFNγ prior to infection where indicated. Detailed isolation and differentiation protocols are described in the Supplementary Methods.

### Infection and functional assays

Macrophages and monocytes were infected with *Mtb* H37Rv::mCherry or wildtype at assay-specific multiplicities of infection. Intracellular bacterial burden was quantified by colony-forming unit (CFU) assays. Nitric oxide production was assessed using the Griess assay, and CXCL10 secretion was measured by ELISA. In selected experiments, the iNOS inhibitor 1400W was used. Full assay protocols are provided in the Supplementary Methods.

### Transcriptomic analyses

Bulk RNA sequencing was performed on IFNγ-treated and *Mtb*-infected macrophages. Libraries were prepared using Smart-seq2 and sequenced on an Illumina NovaSeq platform. Differential gene expression, pathway enrichment, and ortholog analyses were conducted using established bioinformatic pipelines. Public scRNA-seq and bulk RNA-seq datasets were re-analyzed for comparative purposes. Full sequencing and computational details are described in the Supplementary Methods.

### Multiplexed Ion Beam Imaging - Time of Flight (MIBI-TOF) Analysis

Analysis of MIBI-TOF images from non-human primate and human tissue was performed on previously published data sets for the analysis of human granuloma (41) and non-human primates (42). Representative images from granuloma and surrounding tissue were re-analyzed focusing on areas enriched for epithelial cell.

### Statistical analysis

For experimental data, non-parametric statistical tests were used throughout. Differential gene expression analyses were performed using DESeq2 with false discovery rate correction. Additional details are provided in the Supplementary Methods.

## Results

### Lymphocyte - IFNy axis does not induce *NOS2* expression in human lung myeloid cells *in vivo*

IFNγ activated mouse macrophages upregulate iNOS as a key anti-mycobacterial effector to control intracellular *Mtb* growth (43). Transcriptomics allows to characterize potent iNOS^+^ macrophage subsets at single cell resolution (44). To define NOS2 expression in myeloid cells *in vivo*, we reanalyzed seven published scRNA-seq and bulk RNA-seq datasets from humans, nonhuman primates (NHPs), and mice with *Mtb* infection (Table S1). The data set by Wang *et al*. 2023 (45) included lung resections from TB patients with high and low 18F-FDG uptake, reflecting regions of differential inflammation and bacterial burden, while Reichmann *et al.* analyzed bulk RNA-seq from human TB granulomas isolated from lymph nodes (46). In both datasets, *IFNG*-expressing T cells and *CXCL10*⁺ myeloid cells were readily detected, whereas NOS2 transcripts were largely absent from all myeloid clusters (Fig. 1A; Fig. S1A–D). Similar results were observed in NHP granuloma datasets (47, 48), in which NOS2 RNA was undetectable despite the presence of *IFNG*⁺ lymphocytes and *CXCL10*⁺ macrophages (Fig. 1D–F; Fig. S1E–G). In contrast, multiple mouse datasets showed robust *Nos2* expression in *Cxcl10*⁺ myeloid cells in the presence of *Ifng*⁺ lymphocytes (Fig. 1G–I; Fig. S2A–E). Thus, while scRNAseq datasets from mouse allow detailed analysis of *Nos2* expressing myeloid subsets comparable datasets from primates do not contain such *NOS2^+^* myeloid cells *in vivo*.

**Fig. 1.**
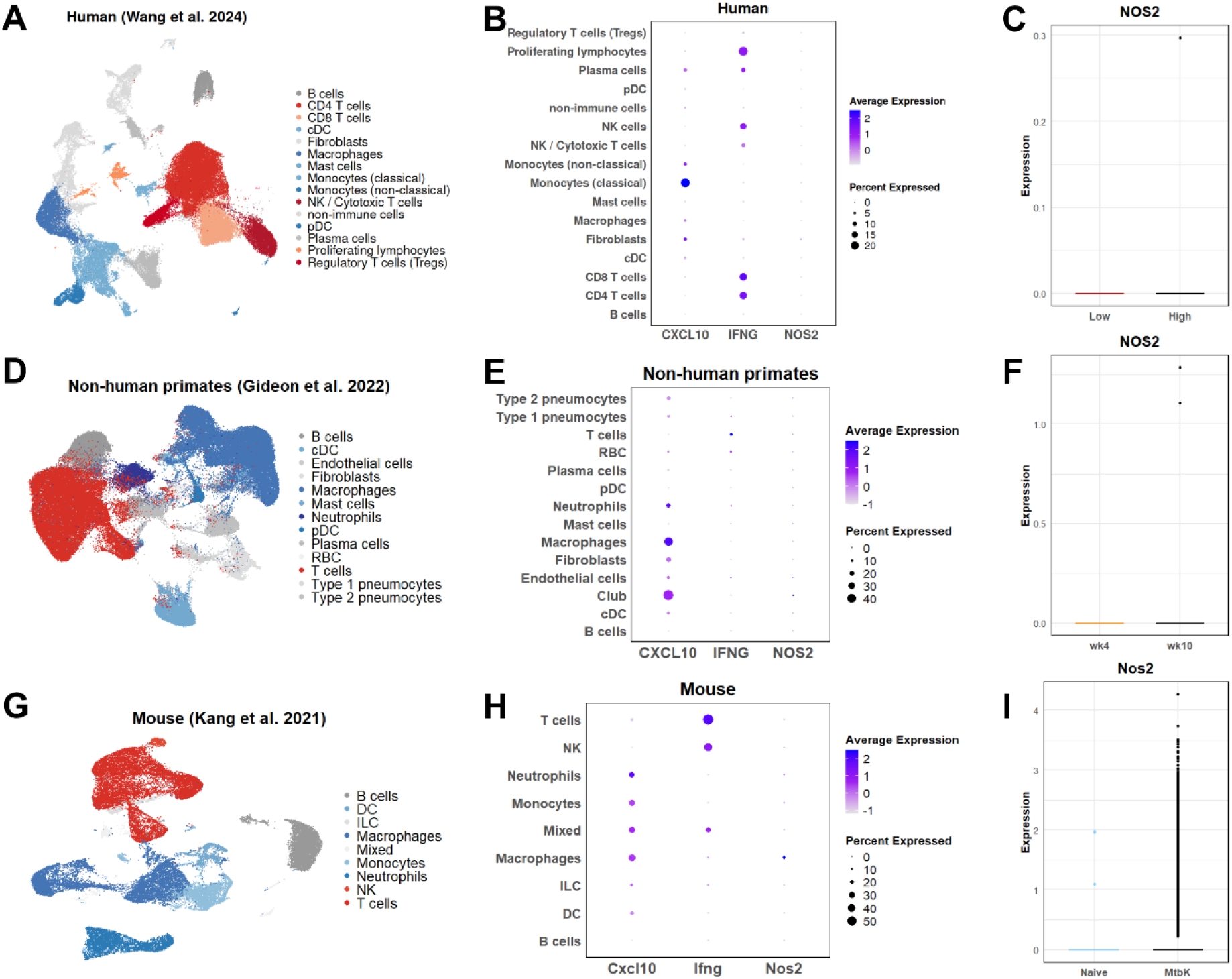
Publicly available scRNAseq data confirm absence of *NOS2* in myeloid cells from human and non-human primate TB granulomas. (A–C) Reanalysis of scRNA-seq data from human lung resections from TB patients (Wang et al., 2024; GSE192483). (A) UMAP showing immune cell clustering. (B) Feature dot plots showing expression of *IFNG*, *CXCL10*, and *NOS2* across immune cell clusters. (C) Box plot of *NOS2* expression in samples from high vs. low 18F-FDG uptake regions (reflecting inflammation level). (D–F) scRNA-seq from TB granulomas of non-human primates (Gideon et al., 2022; GSE200151). (D) UMAP showing immune cell clustering. (E) Feature dot plot showing *CXCL10⁺* myeloid and *IFNG*⁺ lymphocyte populations in the absence of *NOS2* expression. (F) Box plot comparing *NOS2* expression in granulomas from 4 weeks vs. 10 weeks post-infection. (G–I) Mouse lung transcriptomics from *Mtb*-infected mice (Kang et al., 2021; GSE167650). (G) UMAP showing immune cell clustering. (H) Feature dot plot showing robust *Nos2* expression in *Cxcl10*⁺ myeloid clusters in the presence of *Ifng*+ lymphocytes. (I) Box plot comparing *Nos2* expression in naïve vs. *Mtb*-infected lung samples. Data are shown as normalized expression on a log scale.

### iNOS driven *Mtb*-growth restriction is absent in human monocyte-derived macrophages, alveolar macrophages

Our re-analysis of single cell and bulk transcriptomic data indicated lack of *NOS2* upregulation in IFNγ-activated human macrophages *in vivo*. Next, we confirmed this finding in IFNγ-activated macrophages *in vitro*. Consistent with the *in vivo* transcriptomic findings, IFNγ pretreatment induced *CXCL10* expression in both human and mouse macrophages, whereas *NOS2* mRNA and iNOS protein were detected only in mouse macrophages (Fig. 2A–B; Fig. S3A–B). Correspondingly, nitric oxide production was observed exclusively in mouse macrophages and was abrogated by the iNOS inhibitor 1400W (Fig. 2C), which also reversed IFNγ-mediated restriction of intracellular *Mtb* growth (Fig. 2D; Fig. S3C). In contrast, IFNγ pretreatment did not significantly restrict intracellular *Mtb* growth in human macrophages, irrespective of differentiation protocol (Fig. S3D–E), timing of IFNγ exposure (Fig. S4A), or extended pretreatment duration (Fig. S4B), consistent with previous reports (27). Supplementation with the nitric oxide donor DETA-NONOate reduced *Mtb* growth in human macrophages without affecting viability (Fig. S4C–D), indicating that nitric oxide is sufficient to restrict *Mtb* growth but is not endogenously produced. Analysis of primary alveolar macrophages yielded similar results. Human alveolar macrophages upregulated *CXCL10* following IFNγ stimulation (Fig. 2E), whereas only mouse alveolar macrophages displayed detectable iNOS activity after *Mtb* exposure (Fig. 2F). Thus, *in vitro* primary alveolar macrophages and monocyte-derived macrophages mirrored the *in vivo* reanalysis and confirmed the lack of iNOS expression and activity in human macrophages.

**Fig. 2.**
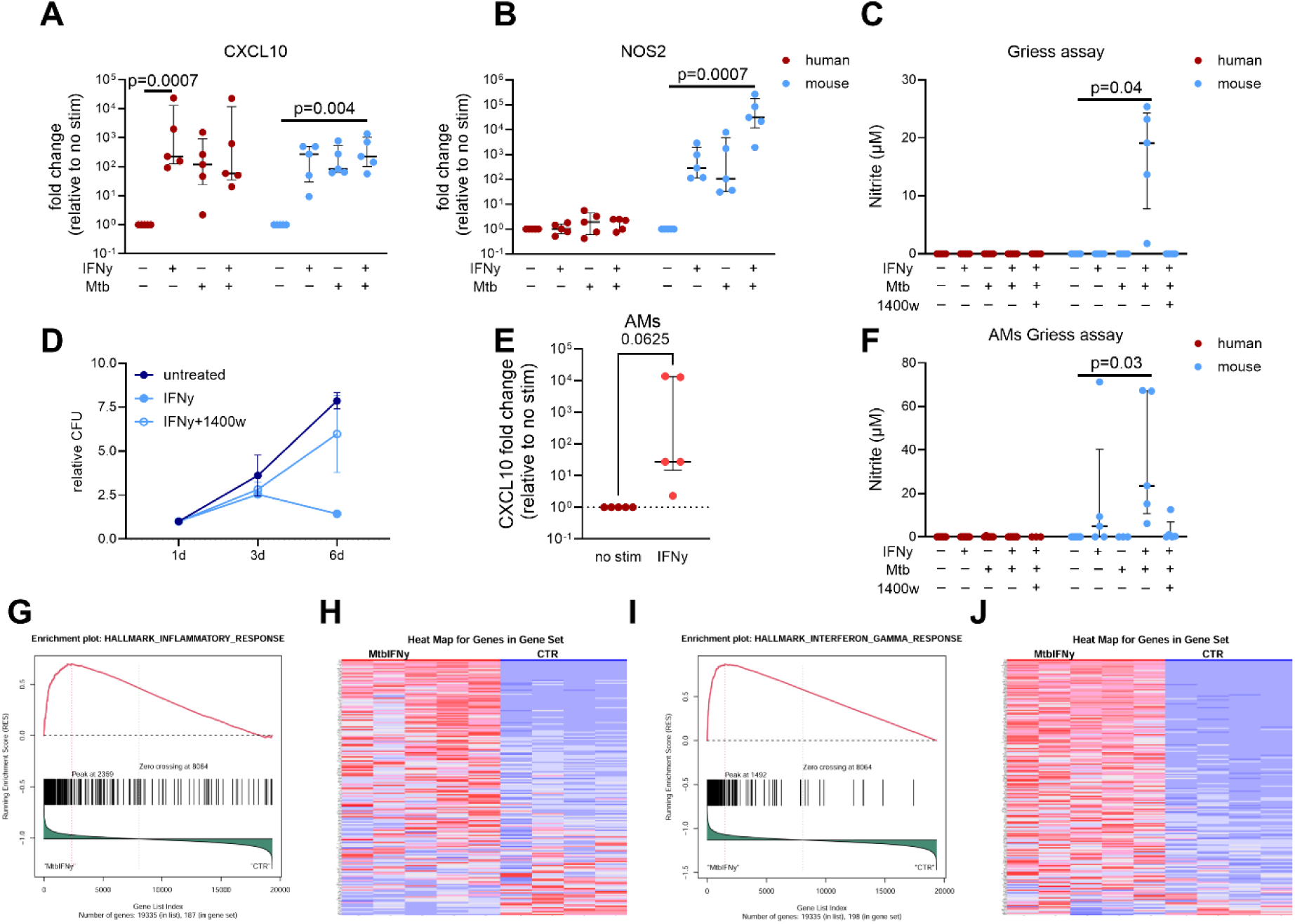
Absence of *NOS2* transcripts and iNOS activity in human myeloid cells besides strong inflammatory responses. (A, B) Human (red; n=5) and mouse (blue; n=5) macrophages were pre-treated with species-specific IFNγ (24 h), followed by infection with *Mtb* H37Rv (MOI 1–2, 24 h). Control samples were untreated or treated with IFNγ or *Mtb* alone. (A-B) *CXCL10* and *NOS2* mRNA expression was determined by qRT-PCR. (C) Nitrite concentrations (iNOS activity) in supernatants were measured by Griess assay (n = 5 per group), where indicated cells were treated with the iNOS inhibitor 1400w after infection (D) Relative *Mtb* CFU in mouse macrophages pre-treated with IFNγ and treated with or without iNOS inhibitor 1400W after infection with *Mtb* H37Rv at a MOI of 0.1-0.2 (n = 2). (E-F) IFNγ response in alveolar macrophages (AMs) from humans (red; n = 5) and mice (blue; n = 5), assessed by *CXCL10* expression (human) and Griess assay (human and mouse). Data shown as median ± interquartile range and individual data points. (A-C, F) *P*-values were determined using Friedman test followed by Dunn’s post hoc correction for multiple comparisons or (E) Wilcoxon rank-sum test.(G-J) Human macrophages were stimulated with IFNγ for 24 h, followed by *Mtb* infection (MOI 1–2) for 24 h. RNA was extracted and sequenced (Smart-seq2). Gene Set Enrichment Analysis (GSEA) for inflammatory and Interferon γ pathway enriched genes of DEGs from IFNγ+*Mtb* vs. control in human macrophages, including enrichment plots and corresponding heatmaps. False discovery rates (FDRs) are indicated in each panel. Statistical analysis was performed using DESeq2 and clusterProfiler.

### IFNγ activates a pro-inflammatory program in human macrophages

To further characterize IFNγ responses, we performed bulk RNA sequencing of IFNγ-treated human or mouse macrophages infected with *Mtb*. Principal component analysis demonstrated clear separation of samples by treatment condition in both species (Fig. S5A–D). Gene set enrichment analysis revealed strong enrichment of IFNγ and inflammatory response pathways in both human and mouse macrophages (Fig. 2G–J; Fig. S5E–H). Orthologous gene expression analysis identified predominantly species-specific responses, with none of the top 50 IFNγ-induced mouse genes significantly upregulated in human macrophages (Fig. S6A–C). Among 15 orthologous genes selectively upregulated in mice, *NOS2, ARG1,* and *ASS1*—enzymes involved in nitric oxide and citrulline metabolism—were undetectable or minimally expressed in human macrophages (Fig. 6D) (49, 50). Consistently, comparison with a published human iNOS-induced gene signature (*51*) revealed positive enrichment in mouse macrophages but negative enrichment in human macrophages (Fig. S6E). Together, these results implicate a robust IFNγ mediated priming of human macrophages with *NOS2* and iNOS activity as key contributors to species-specific differences in macrophage responses to *Mtb*.

### Lack of IFNγ-induced iNOS activity is commonly observed in mammalian myeloid cells

Mouse but not human IFNγ-primed macrophages show high level of iNOS activity after stimulation with the TLR4 ligand LPS (52). To test whether the lack of iNOS activity is specific to human myeloid cells or more universally absent in larger mammalian species, we extended the analysis to monocytes from six additional species: rhesus macaque (RM), cynomolgus macaque (CM), goat, sheep, pig, and cattle (Fig. 3A). These species are natural hosts for TB-causing mycobacteria. Monocytes from all species upregulated *CXCL10* after species-specific IFNγ stimulation (Fig. 4B). *NOS2* expression was moderately induced in goat, sheep, and cattle after IFNγ treatment (Fig. 4C), but only cattle and mouse monocytes showed detectable iNOS activity after IFNγ+LPS treatment (Fig. 4D). Thus, from all 8 mammalian species (incl. mouse and human) only mouse and cattle myeloid cells showed detectable iNOS activity. Taken together, these findings suggest that IFNγ induced *NOS2* expression and iNOS activity is largely absent in myeloid cells from humans and most other mammals, suggesting Th1 mediated control of intracellular growth of *Mtb* via iNOS activity in mouse myeloid cells is rather uncommon in other mammalian species.

**Fig. 3.**
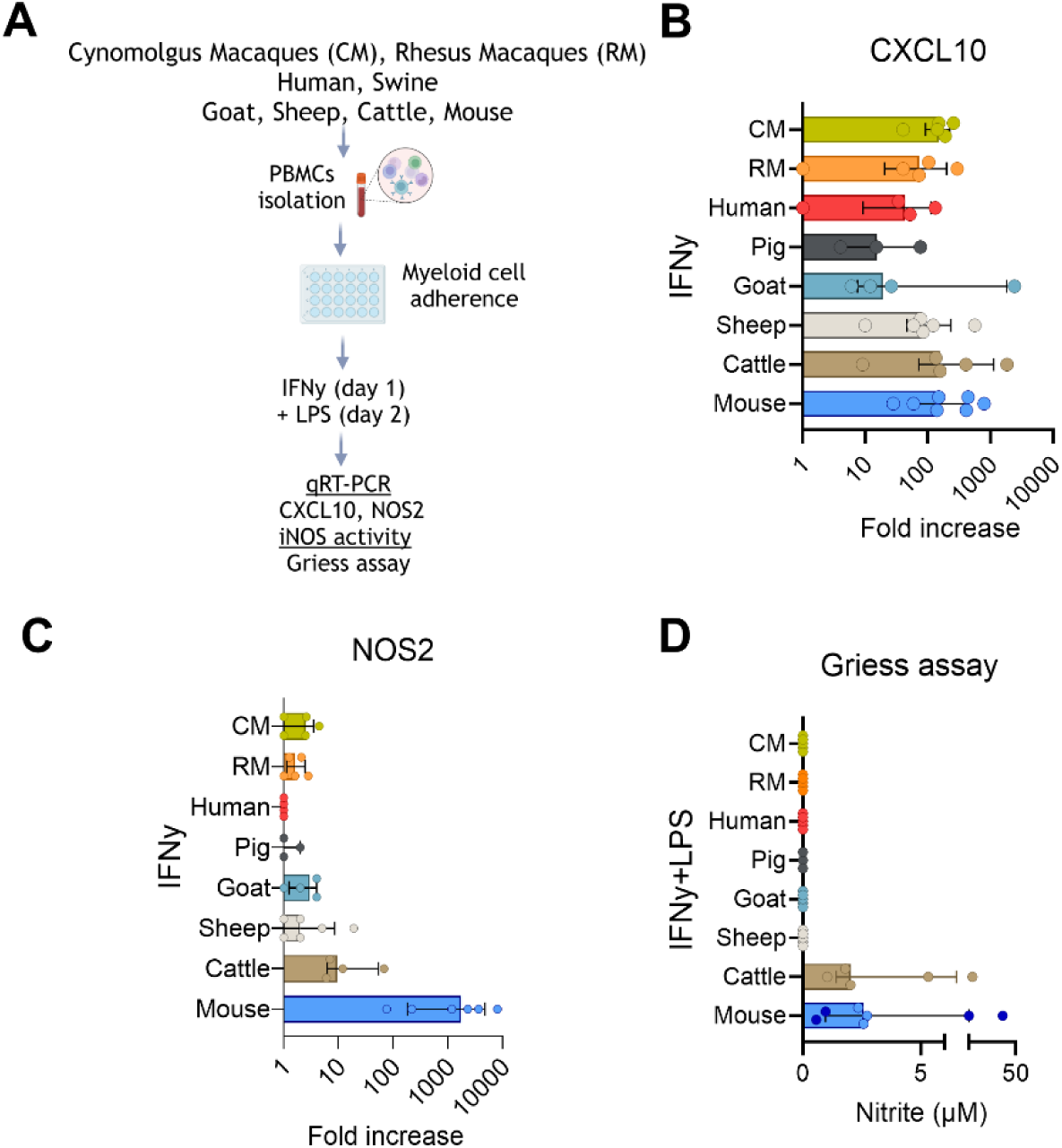
Lack of iNOS expression is common in myeloid cells from different mammalian species. (A-D) Cross-species comparison of monocytes (n = 3–5 per species) from NHPs (cynomolgus macaques = green, rhesus macaques = orange, human = red, pig = dark grey, goat = turquoise, sheep = light grey, cattle = brown and mouse = blue (dark blue = monocytes isolated from blood sample for Griess assay; light blue = monocytes isolated from bone marrow) stimulated with species-specific IFNγ ± 1µg/ml LPS. (H) *CXCL10* induction, (I) *NOS2* expression, and (J) iNOS activity. Data shown as median ± interquartile range and individual data points.

**Fig. 4.**
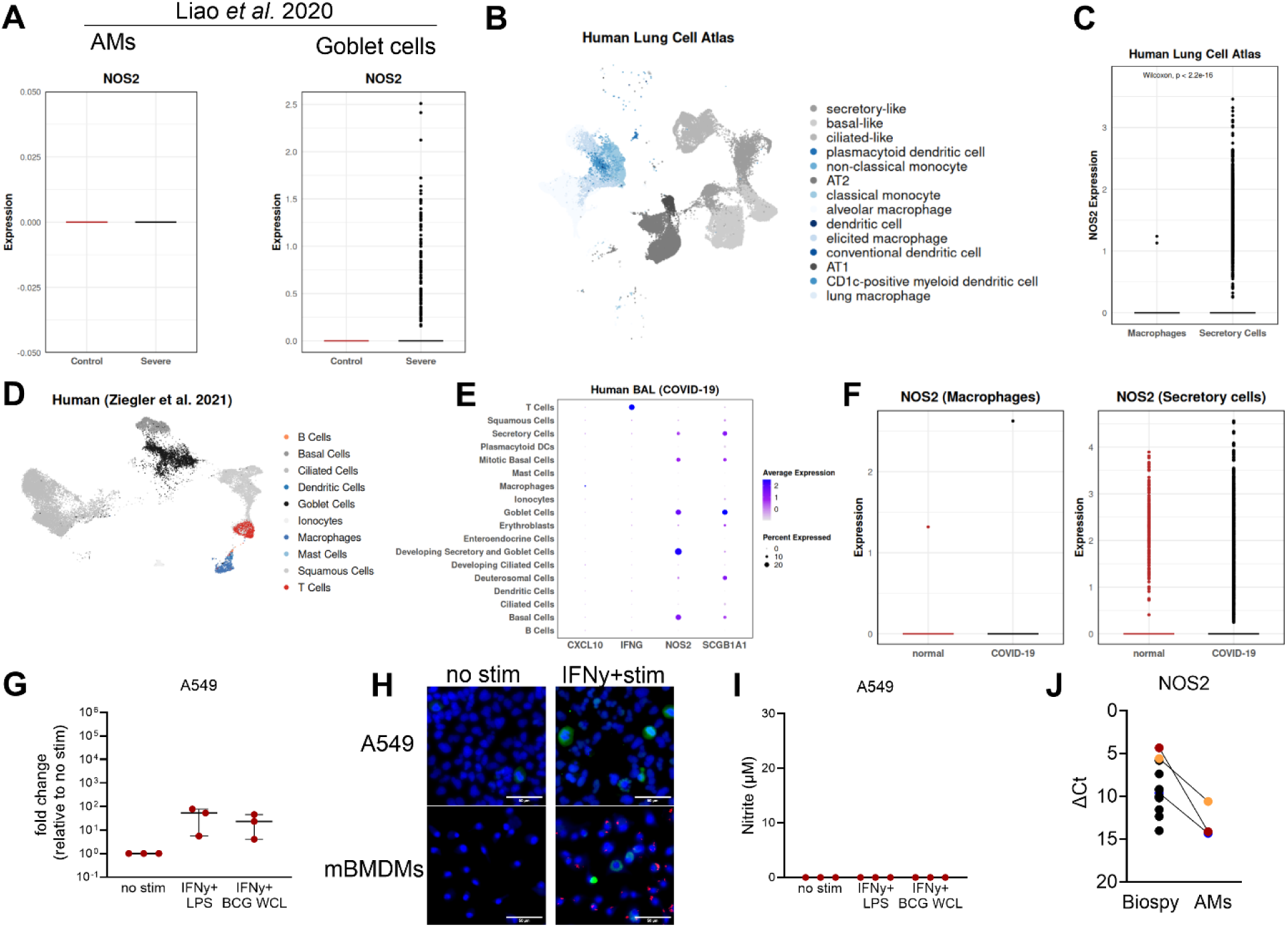
Human respiratory epithelial cells express *NOS2* at steady state and during inflammation. (A) Box plots of *NOS2* expression in alveolar macrophages (AMs) and goblet cells from scRNAseq of BALF in COVID-19 (Liao et al., 2020; GSE145926). (B–C) Human Lung Cell Atlas (https://data.humancellatlas.org/hca-bio-networks/lung/atlases/lung-v1-0). (B) UMAP of myeloid (shades of blue) and epithelial cells (shades of grey and secretory cells in black) and (C) box plot of *NOS2* expression in secretory cells vs. macrophages, Wilcoxon p < 2.2e–16. (D–F) scRNAseq of nasopharyngeal swabs (Ziegler et al., 2021; SCP1289). (D) UMAP of, lymphocytes (shades of red), myeloid (shades of blue) and epithelial cells (shades of grey and secretory cells in black). (E) Feature dot plot of *CXCL10*, *IFNG*, *NOS2*, *SCGB1A1*. (F) Box plots for *NOS2* expression by cell type and condition (F). (G) *NOS2* RNA expression in A549 cells (qRT-PCR) stimulated with IFNγ, LPS, or BCG WCL (n=3 independent experiments). (H) iNOS protein staining in A549 cells and mouse BMDMs after IFNγ + stim (A549 = IFNγ + BCG WCL, mBMDMs (IFNγ + Mtb H37Rv mcherry), scale bar= 20 µm; blue = nuclei DAPI staining, green = iNOS, red = *Mtb* mcherry. (I) Nitrite levels in A549 supernatants after stimulation (n=3 independent experiments). (J) ΔCt values (reversed y-axis) for *NOS2* RNA in human upper airway biopsies vs. AMs (biopsies n=13; AMs n = 3 paired samples indicted by color code). *Ρ*-values were determined using two-sided Wilcoxon rank-sum tests (panel C: P < 2.2 × 10⁻¹⁶; panel J: P = 0.25). Other panels are descriptive with biological replicates.

### Human airway secretory epithelial cells express *NOS2*

In contrast to our findings, nitric oxide production and iNOS activity has been reported in human TB and other respiratory conditions, proposing that other cell types than macrophages might be responsible for this activity (33, 34). Therefore, we examined whether non-myeloid cell types contribute to NOS2 expression. Reanalysis of scRNA-seq data from bronchoalveolar lavage samples of COVID-19 patients (53) revealed enrichment of *NOS2*⁺ cells within secretory epithelial clusters compared with alveolar macrophages (Fig. 4A). This pattern was confirmed in the Human Lung Cell Atlas (54) (normal respiratory sample data sets) where *NOS2* expression was most prominent in secretory epithelial cells of the upper respiratory tract (Fig. 4B–C; Fig. S7). Similar *NOS2* and *SCGB1A1* co-expression was observed in nasopharyngeal epithelial cells from COVID-19 patients (55) (Fig. 4D–F). In contrast, NOS2 expression was not detected in epithelial cells from murine nasal mucosa following influenza infection or IFNα treatment (56, 57) (Fig. S8). *In vitro* stimulation of human A549 type II pneumocyte epithelial cells with IFNγ and LPS or BCG whole cell lysate (WCL) induced *NOS2* mRNA and iNOS protein, but not measurable iNOS activity (Fig. 4G–I). Finally, we performed qPCR on human airway biopsy samples enriched for bulk expression of *SCGB1A1* and *MUC5AC* (Fig. S9A) and found *NOS2* relative expression elevated in human biopsy samples from the upper airway compared to alveolar macrophages (Fig. 4J). In conclusion, reanalysis of publicly available scRNAseq data sets enriched for secretory epithelial cells highly expressed *NOS2* RNA proposing this cell type as a relevance source of iNOS in humans.

### Non-human primate and human TB granuloma are surrounded by iNOS expressing lung epithelial cells

Although *NOS2* expression was evident in human airway epithelium, it was not consistently detected in human (45) or NHP TB granuloma scRNA-seq datasets (56), likely reflecting the paucity of epithelial cells in granuloma-containing tissue sections and/or limited epithelial-cell capture efficiency by scRNA-seq (Fig. 1A; Fig. S9B–D). We therefore sought to evaluate NOS2-positive epithelial cells within intact granuloma-associated tissue architecture using multiplexed image analysis. To this end, we reanalyzed previously published granuloma MIBI-TOF datasets from McCaffrey and colleagues, including both NHP (42) and human (41) granuloma tissue sections (Fig. 5A-B). In this reanalysis, we prioritized sections enriched for epithelial cells adjacent to granulomatous lesions. Epithelial cells were identified using pan-cytokeratin (PanCK), with vimentin (VIM) included as a broad cytoplasmic marker. CD31 labeled endothelial cells, αSMA labeled myofibroblasts in NHP sections (Fig. 5A) and CD45 labeled hematopoietic-derived immune cells in human sections (Fig. 5B). In both species, iNOS staining was detected predominantly outside granulomas and colocalized with PanCK-positive epithelium rather than with CD45-positive immune cells within granulomatous structures. In combination with the histology image, the iNOS staining clearly clustered in an area of lung parenchyma not associated with the macrophage-rich myeloid core where epithelioid macrophages co-localize. Collectively, these findings indicate that iNOS-positive epithelial cells are a prominent feature adjacent to human and NHP TB granulomas.

**Fig. 5.**
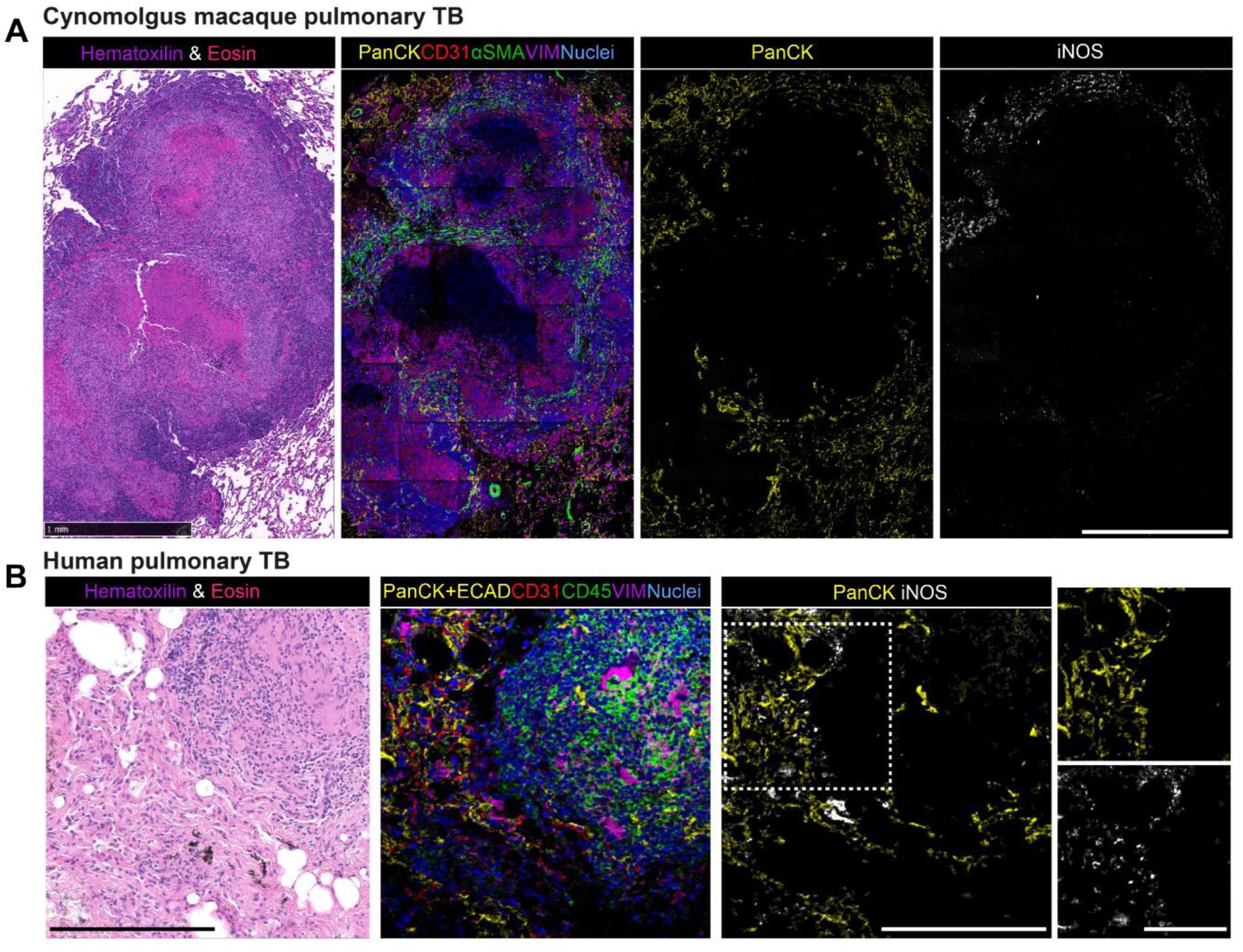
Non-human primate and human epithelial cells express iNOS in close proximity to granuloma structures. (A) Representative images from a cynomolgus macaque pulmonary TB granuloma analyzed by McCaffrey & Delmastro et al 2025. Images show hematoxylin & eosin staining (left) and an adjacent serial tissue section analyzed via MIBI-TOF (middle left, middle right, right) for pan-cytokeratin (PanCK, yellow), CD31 (red), alpha-smooth muscle actin (aSMA, green), vimentin (VIM, magenta), histone H3 (nuclei, blue), and iNOS (white). The iNOS and panCK stains were performed on adjacent serial sections. Scale bars represent 1 mm. (B) Representative images from a human pulmonary TB granuloma analyzed by McCaffrey et al 2022. Images show hematoxylin & eosin staining (left) and an adjacent serial tissue section analyzed via MIBI-TOF (middle left, middle right, right) for pan-cytokeratin and E-cadherin (PanCK+ECAD, yellow), CD31 (red), CD45 (aSMA, green), vimentin (VIM, magenta), dsDNA (nuclei, blue), and iNOS (white). The iNOS and panCK stain were performed on the same sections. Scale bars represent 200 µm for larger images and 100 um for zoomed insets (right).

In conclusion, our findings demonstrate that IFNγ activation fails to induce a functional anti-mycobacterial program in human macrophages, in stark contrast to mouse macrophages. In particular, we identified *NOS2*/iNOS expression and activity as a key species-specific effector absent in human and most non-murine myeloid cells. Reanalysis of *in vivo* datasets confirmed this pattern and revealed secretory epithelial cells in the human upper airway and epithelial cells in close proximity to TB granuloma as an alternative source of iNOS in primates. These results emphasize fundamental interspecies differences in IFNγ-driven host defense and caution against direct extrapolation from mouse to human immune responses.

## Discussion

In this study, we used a stepwise, cross-species approach to define the cellular sources and functional relevance of iNOS during *Mtb* infection. Reanalysis of publicly available scRNA-seq datasets from human and nonhuman primate TB lesions revealed a consistent absence of *NOS2* expression in myeloid cells despite robust IFNγ-associated inflammatory signatures, in striking contrast to murine granulomas where *Nos2* expression was readily detected. In light of these *in vivo* observations, we assessed IFNγ-induced macrophage responses *in vitro* and found that, although IFNγ robustly activated inflammatory programs in human macrophages, it failed to induce *NOS2* transcription, iNOS activity, or restriction of intracellular *Mtb* growth. Extending this analysis across species demonstrated that this lack of IFNγ-induced iNOS activity is conserved across multiple mammalian myeloid compartments. In contrast, airway epithelial cells emerged as a prominent source of *NOS2* expression in humans, and multiplexed ion beam imaging spatially localized iNOS protein to epithelial cells adjacent to granulomatous lesions. Together, these data redefine the cellular context of nitric oxide biology in human tuberculosis and challenge IFNγ–iNOS paradigms derived from murine models.

Although IFNγ activated comparable inflammatory pathways across species, control of intracellular *Mtb* growth was observed only in mouse macrophages. Previous studies similarly reported unrestricted *Mtb* growth in IFNγ-stimulated human monocyte-derived and alveolar macrophages under a range of experimental conditions (*25–27, 30*). While restricted *Mtb* growth has been observed in human macrophage cultures under hypoxia or in the presence of additional effector molecules such as TNFα, these effects were not linked to iNOS activity (32). In our and other studies (17), IFNγ-dependent but iNOS-independent effector responses were insufficient to control *Mtb* growth in human macrophages, reinforcing the central role of iNOS in murine—but not human—macrophage antimicrobial activity.

Our findings contrast with reports detecting iNOS protein in human and nonhuman primate tuberculosis lesions and nitric oxide in exhaled breath of TB patients (33, 36, 42, 58). These observations have been interpreted as evidence for macrophage-derived iNOS activity in human tuberculosis. However, the consistent absence of *NOS2* transcripts in macrophages from human and primate scRNA-seq datasets (45, 46) suggests that such cells are either rare, transient, or express NOS2 at levels below detection thresholds. Moreover, exhaled nitric oxide levels in TB patients are generally low and do not consistently correlate with disease severity or progression, raising questions about the functional relevance of macrophage-derived nitric oxide in humans (59–61). In contrast, we observed reproducible NOS2 expression in airway epithelial cells across multiple datasets and conditions, consistent with prior reports identifying epithelial iNOS as a major determinant of nitric oxide production in the respiratory tract (62, 63). Our *in vitro* work also confirmed presence and upregulation of *NOS2* RNA in epithelial cells. However, the A549 type II pneumocyte cell line probably does not reflect the phenotype of *NOS2* positive cells as observed *in vivo* (*64*). While NOS2 expression in epithelial cells did not consistently translate into measurable nitric oxide production *in vitro*, spatial proteomic analyses of intact granuloma tissue sections identified epithelial cells—rather than macrophages—as the predominant iNOS^+^ population in proximity to granulomatous lesions. The functional contribution of epithelial-derived nitric oxide to TB immunity remains unclear. Epithelial cells may influence local immune responses, bacterial containment, or tissue pathology rather than directly mediating intracellular bacterial killing. Notably, epithelial *NOS2* expression appeared more prominent in human tissues than in available murine datasets (55, 57), underscoring additional species-specific differences that warrant further investigation.

This study has several limitations. *In vitro* macrophage differentiation and culture conditions may not fully recapitulate *in vivo* phenotypes, and donor numbers for some species were limited by feasibility. Low-level NOS2 expression or nitric oxide production may fall below detection thresholds in certain experimental settings. Additionally, while we focused on *NOS2* as a key IFNγ-induced effector, other species-specific antimicrobial pathways may contribute to *Mtb* control and were not explored in depth here.

Our findings have important implications for TB vaccine development particularly considering the respiratory parenchyma. Many current vaccine strategies aim to elicit strong IFNγ-producing CD4⁺ T cell responses based on murine models in which IFNγ-driven iNOS induction in macrophages is a central mechanism of protection. Our data indicate that, in humans, IFNγ robustly activates inflammatory programs but does not induce comparable bactericidal effector functions in macrophages. These interspecies differences challenge the use of IFNγ responses as universal correlates of protection and highlight the need to reconsider how preclinical efficacy is translated into clinical settings. Future vaccine strategies may need to incorporate alternative or complementary mechanisms of protection, including modulation of epithelial responses, trained immunity, or cytokine networks acting in concert with IFNγ in the absence of iNOS activity. Ultimately, rational TB vaccine design will require a deeper understanding of human-specific immune mechanisms rather than direct extrapolation from murine models.

## Author contributions

Conceptualization: AD, BDB and BC. Methodology: FS, BZ, JP, SS, EFM and BC. Investigation: FS, BZ, JP, BES, EFM and BC. Formal analysis: FS, BZ, JP, SS, EFM and BC. Resources: BS, RW, FV, BES, AD, BDB, EFM and BC. Visualization: FS, BZ, JP, EFM and BC. Supervision: AD and BC. Writing—original draft: BC. Writing—review & editing: FS, BZ, JP, SS, BS, RW, FV, BES, AD, BDB, EFM, BC

## Acknowledgments

We acknowledge Laura Timm, Gabriele Stooß and Ulrike Zedler for the outstanding technical support. We are grateful to Bärbel Hammerschmidt, Yolanda Marschner and Dr. Charlotte Schröder for their support and providing animal samples. We would like to thank the members of the University Hospital Greifswald transfusion medicine and pneumology for providing human samples. The *Mtb* reporter strain was generously provided by Stefan H.E. Kaufmann.

