## Supplemental Material for "Species-specific iNOS expression distinguishes epithelial and myeloid IFNγ responses in tuberculosis"

**Affiliations**

**Detailed Material and Methods**

### Ethical Approval

All biological sample collections were conducted under appropriate ethical oversight and in accordance with institutional and national guidelines. Human PBMCs were isolated from buffy coats obtained from anonymous healthy blood donors through the blood donation center of the University Hospital Greifswald. Donors provided written informed consent, and the use of these samples for research was approved by the Ethics Committee of the University Hospital Greifswald as part of the protocol from the Institute for Transfusion Medicine (project leaders: Prof. Dr. Greinacher and Prof. Dr. Thiele; approval ID BB 014/14). For bronchoalveolar lavage (BAL) fluid and airway biopsy samples, written informed consent was obtained prior to collection. Samples were derived from excess clinical material collected during routine diagnostic procedures. Patients with diagnosed lung infections were excluded. This study was approved by the Ethics Committee of the University Hospital Greifswald (approval ID BB 058/19). EDTA blood samples from healthy cattle, swine, goats, and sheep were collected at the Friedrich-Loeffler-Institut (FLI) under minimal stress conditions and without euthanasia. These procedures were approved by the local authority (Landesamt für Landwirtschaft, Lebensmittelsicherheit und Fischerei Mecklenburg-Vorpommern) under protocol numbers LALLF 7221.3-2-041/17 and 7221.3-2-001/23. Mouse samples (bone marrow, blood, and BAL fluid) were collected post-mortem from animals euthanized without prior experimental procedures, in accordance with §4 Absatz 3 of the German Animal Welfare Act ("Töten zur Organentnahme"). NHPs PBMCs in this study were obtained from tissues repositories of the Biomedical Primate Research Centre, Rijswijk, Netherlands. The PBMCs were banked as surplus either from prior ethically approved NHP research protocols or regular health check procedures. Ethical approval for collecting blood samples at the time was provided by the institutional ethics committee (Dierexperimentencommissie, DEC). Samples were selected from untreated control conditions and by the availability of at least $50x10^6$ cells of any particular individual.

**PBMCs isolation**

Human buffy coats were diluted 1:2 with D-PBS (VWR) and subjected to density gradient centrifugation using Pancoll (ρ = 1.077 g/ml; PAN-Biotech) at room temperature (RT) for 30 minutes at 400 × g, without brake. The peripheral blood mononuclear cell (PBMC) layer was collected and washed twice with RPMI 1640 (Biowest). Cell debris and platelets were removed by low-speed centrifugation at 100 × g for 20 minutes. Residual erythrocytes were lysed in 1 ml red blood cell (RBC) lysis buffer (BioLegend) for 5 minutes at 4°C. The reaction was quenched by adding RPMI 1640 supplemented with 10% fetal calf serum (FCS; PAN-Biotech). Viable PBMCs were counted using trypan blue exclusion with either a Neubauer counting chamber or a Bio-Rad TC20 automated cell counter. For livestock species (cattle, sheep, goat, and swine), blood was collected in EDTA tubes (Sarstedt) and processed as described for human PBMCs, except that density centrifugation was performed at 800 × g. PBMCs from these species were used fresh, without cryopreservation. For non-human primates (NHPs), PBMCs were isolated from heparinized blood by density gradient centrifugation. Lymphoprep (Axis-Shield, UK) was used for rhesus macaques, and Percoll (GE Healthcare, IL, USA) was used for cynomolgus macaques. Cells were resuspended in RPMI medium supplemented with 50% FCS and 10% dimethyl sulfoxide (DMSO) and stored long-term in liquid nitrogen.

**Differentiation of macrophages**

Human monocytes were isolated from PBMCs by magnetic cell sorting at 4°C. PBMCs were incubated with biotin-conjugated anti-human CD14 antibody (clone M5E2, BioLegend) for 15 minutes, washed with MACS buffer (0.5% bovine serum albumin [BSA, Sigma], 2 mM EDTA [Invitrogen] in D-PBS), and subsequently incubated with streptavidin-coated nanobeads (MojoSort™, BioLegend) for 15 minutes. After a final wash, the cell suspension was transferred to a 5 ml polypropylene tube and was placed into a magnet (StemCell) for positive selection of $CD14^+$ monocytes. Following several rounds of magnetic purification and a washing step, monocytes were plated at a density of $0.1 \times 10^6$ viable cells/well in 96-well plates or $0.35 \times 10^6$ cells/well in 48-well plates in RPMI 1640 supplemented with 10% heat-inactivated human serum (PAN-Biotech), 2 mM L-glutamine, and 1 mM sodium pyruvate. Where indicated, human serum was replaced with 10% FCS and either 50 ng/ml recombinant human GM-CSF or M-CSF (both from BioLegend). Half of the medium was replaced every 2–3 days. Cells were used for experiments after 6–10 days of differentiation at 37°C and 5% $CO_2$. Mouse macrophages were generated from bone marrow of C57BL/6 mice. Bone marrow cells were flushed with D-PBS, washed, and resuspended in high-glucose DMEM (Fisher Scientific) supplemented with 20% L929 cell-conditioned medium, 10% FCS, 5% horse serum, 2 mM L-glutamine, and 1 mM sodium pyruvate. Cells were cultured in petri dishes (Sarstedt) at 37°C and 5% $CO_2$ for 7 days, with half of the medium replaced on day 4. Unless stated otherwise, fully differentiated macrophages from human and mouse origin were maintained in RPMI 1640 with 10% FCS, 2 mM L-glutamine, and 1 mM sodium pyruvate. Where indicated, cells were stimulated with 500 ng/ml species-specific recombinant IFNγ (Peprotech).

**A549 epithelial cell culture and stimulation**

The human alveolar epithelial cell line A549 (FLI, cell bank) was maintained in RPMI 1640 supplemented with 10% fetal calf serum, 2 mM L-glutamine, and 1 mM sodium pyruvate, identical to the culture medium used for mature macrophages. Cells were seeded at a final density of $0.5 \times 10^6$ cells per well in 24-well plates or $0.1 \times 10^6$ cells per well in 96-well plates. For stimulation experiments, cells were treated with 500 ng/ml recombinant human IFNγ (Peprotech) 24 h prior to stimulation with 1 µg/ml LPS (InvivoGen) or with 10 µg/ml *Mycobacterium bovis* BCG WCL, as indicated in Fig. 6. Cells were harvested for RNA extraction, protein staining, or nitrite quantification in supernatants after 24 hours of stimulation.

**Isolation of alveolar macrophages and airway biopsy samples**

Human bronchoalveolar lavage fluid (BALF; 10–20 ml in sterile saline) and airway pinch biopsies were obtained from the University Hospital Greifswald and processed within 1–3 hours storage at 4°C. We included a total of 18 samples (male=10 median age 60 and range of 32-70, female=8 median age of 55 and range of 34 – 71). The main indication for bronchoscopy in these patients was the diagnostic evaluation of suspected sarcoidosis or other inflammatory lung diseases. Samples for each assay were used based on availability and total cell numbers. BALF was filtered through a 70 µm cell strainer (VWR), centrifuged at 400 × g for 10 minutes, and adherent alveolar macrophages (AMs) were enriched by plating and washing off non-adherent cells after 30 minutes at 37°C and 5% $CO_2$. Cells were cultured in RPMI 1640 supplemented with 10% FCS, 2 mM L-glutamine, 1 mM sodium pyruvate, and penicillin/streptomycin (Gibco). Mouse alveolar macrophages were isolated post-mortem by flushing the lungs via the trachea with 3 × 1 ml of ice-cold D-PBS containing 0.1% EDTA and penicillin/streptomycin. The collected BALF was processed and cultured as described for human samples.

**Monocyte multi-species comparison assay**

PBMCs from humans, cattle, sheep, goats, swine, and non-human primates (NHPs) were plated at $1 \times 10^6$ cells/well (96-well) or $10 \times 10^6$ cells/well (48-well). Freshly isolated PBMCs were used for all species except NHPs, which were thawed from cryopreserved aliquots. After 2 hours of incubation at 37°C and 5% $CO_2$, non-adherent cells were removed, and adherent monocytes were cultured in RPMI 1640 with 10% FCS, 2 mM L-glutamine, 1 mM sodium pyruvate, and penicillin/streptomycin. Mouse monocytes were isolated either from bone marrow using the MojoSort™ mouse monocyte isolation kit (BioLegend) or from blood. For blood, leukocytes were collected from the abdominal vein into heparin tubes (Sigma), and red blood cells were lysed using RBC lysis buffer for 15 minutes at room temperature. Cells were washed in RPMI 1640 + 10% FCS and plated at $1 \times 10^6$ leukocytes per well in 96-well plates. After 2 hours, non-adherent cells were washed off with D-PBS and adherent monocytes were cultured under the same conditions as other species. Where indicated, wells were treated immediately after plating with 500 ng/ml of species-specific recombinant IFNγ (KingFisher for livestock; Peprotech for human, mouse, and NHP; NHPs PBMCs were treated with confirmed cross-reactive human rec. IFNγ). After 24 hours, cells were stimulated with 1 µg/ml LPS (E. coli K12, InvivoGen) or left untreated. Supernatants were collected for Griess assay, and cells in 96-well plates were stained with DAPI for cell counting by microscopy. Cells in 48-well plates were lysed in 1 ml Trizol™ (Invitrogen) after 24 hours and frozen at –80°C for RNA extraction.

**Bacterial cultures**

*Mycobacterium tuberculosis* H37Rv::mCherry or wildtype were used for all experiments. The mCherry reporter strain was generously provided by Stefan H.E. Kaufmann. Bacteria were cultured to mid-log phase at 37°C with shaking at 110 rpm in Middlebrook 7H9 broth supplemented with 10% ADC, 0.2% glycerol, and 0.05% Tween-80.

**Infection of myeloid cells with *Mtb***

To prepare the *Mtb* stock, 1 ml of culture was centrifuged at 2000 × g for 5 minutes, and the pellet was resuspended in 1 ml D-PBS. Optical density (OD) was measured to estimate bacterial concentration. Single-cell suspensions were prepared using a syringe and diluted in RPMI 1640 supplemented with 10% FCS, 2 mM L-glutamine, and 1 mM sodium pyruvate. Multiplicity of infection (MOI) was adjusted depending on the assay: MOI 0.1–0.2 for CFU, MOI 1–2 for Griess assay and RNA extraction. After replacing medium with bacteria-containing suspension, cells were incubated at 37°C and 5% $CO_2$. For CFU assays, non-phagocytosed bacteria were removed by washing with 200 µl warm D-PBS after 60–120 minutes. Medium was replaced with fresh RPMI. Where indicated, 100 µM of the iNOS inhibitor 1400W hydrochloride (Biomol) was added either at infection (Griess, RNA) or after washing steps (CFU).

**Colony forming units (CFU)**

At 1 (60-120min after exposure and washing step), 3, or 6 days post-infection, cells were lysed in 100 µl of 0.01% Triton X-100 in PBS for 1–5 minutes. Lysates were serially diluted (1:10–

1:1000) in D-PBS with 0.1% Tween-80. Aliquots (50 µl) of the 1:100 and 1:1000 dilutions were plated on Middlebrook 7H11 agar. Plates were incubated at 37°C and 5% $CO_2$ for at least 3 weeks before colony enumeration.

**Quantitative reverse transcriptase polymerase chain reaction (q-RT PCR)**

RNA was extracted from Trizol™ samples using PhaseMaker tubes and RNase-free chloroform (at a ratio of 1:5). After mixing and a 3-minute incubation, samples were centrifuged at 12,000 × g for 15 minutes at 4°C. The aqueous phase was transferred to RNase-free tubes, and RNA was precipitated with 20 µg glycogen and 200 µl cold isopropanol for 60 minutes at –20°C, followed by centrifugation at 12,000 × g for 15 minutes. Pellets were washed with 75% ethanol, centrifuged again, and residual liquid was evaporated. RNA pellets were dissolved in 50 µl nuclease-free $H_2O$ and heated to 60°C for 15 minutes. cDNA was synthesized from 100 ng/µl RNA using the High-Capacity cDNA Reverse Transcription Kit (Applied Biosystems™) following the manufacturer's instructions. qPCR reactions contained 2 µl of cDNA, 10 µM forward and reverse primers, and PowerUp™ SYBR Green Master Mix (Applied Biosystems™) in a total volume of 10 µl. Reactions were run on a QuantStudio6 qPCR system. Primer sequences and PCR cycling conditions are listed in Tables S2 and S3. Primers were designed using NCBI Primer-BLAST, with the exception of published primers for SCGB1A1 and MUC5AC. All primers spanned at least one exon-exon junction and were validated by melting curve, gel electrophoresis, Sanger sequencing, and amplification efficiency testing (90–110%). For *NOS2*, due to low expression in human samples, one primer pair was designed within a single exon, and a second confirmatory pair spanned multiple exons.

**Griess assay**

Supernatants from myeloid cell cultures were used for Griess assay to assess nitric oxide (NO) production, an indirect measure of iNOS activity. For each sample, 100 µl of supernatant was mixed with 100 µl Griess reagent (4% w/v in distilled $H_2O$) in an ELISA plate, in duplicate. Plates were incubated in the dark for 20 minutes, and absorbance was measured at 540 nm using a TECAN plate reader. Nitrate concentration was calculated using a sodium nitrate standard curve (6.25–400 µM).

**Fluorescent microscopy**

To quantify viable macrophages, cells were stained with a green fluorescent viability dye for 15 minutes at 37°C and 5% $CO_2$. After washing with D-PBS, cells were fixed with 4% paraformaldehyde for 10 minutes at room temperature and stained with DAPI for 5 minutes, followed by overnight fixation in 4%PFA. Imaging and high-throughput quantification were performed using the CellInsight™ CX7 High Content Analysis Platform (ThermoFisher).

**BulkRNAseq**

For transcriptomic analysis, macrophages were differentiated in 48-well plates at a density of 1 × $10^6$ cells per well. Cells were pre-treated with 500 ng/ml recombinant IFNγ for 24 hours, followed by infection with *Mtb* at a multiplicity of infection (MOI) of 1 for an additional 24 hours. Control conditions included untreated cells, IFNγ treatment alone, or *Mtb* infection alone. RNA was extracted as described for qRT-PCR, and RNA quality was assessed using an Agilent 2100 Bioanalyzer. Only samples with a RNA integrity number (RIN) > 7 were included for library preparation.

**Library preparation**

Total RNA was purified using RNA-SPRI beads and ethanol washes. Whole-transcriptome amplification (WTA) was performed using the Smart-seq2 protocol (1). Briefly, RNA was mixed with RNA-SPRI beads at a 1:2.2 ratio, incubated, washed in 80% ethanol, and dried prior to reverse transcription. First- and second-strand cDNA synthesis was carried out using master mixes and thermocycler programs described in Table S4. Amplified cDNA was purified with DNA-SPRI beads and 70% ethanol washes, then eluted in TE buffer. cDNA quality and quantity were assessed using a Qubit fluorometer and Bioanalyzer. Nextera XT was used for tagmentation and indexed adapter ligation, followed by PCR amplification. Final libraries were cleaned with DNA-SPRI beads, eluted in TE buffer, and stored at –20 °C until sequencing.

**Sequencing and analysis**

A total of 38 RNA libraries (human, n = 20; mouse, n = 18) were sequenced on an Illumina NovaSeq platform (Technical University Dresden) using 2 × 100 bp paired-end reads, with ~20 million reads per library. FASTQ files were aligned to the reference genomes (Homo sapiens, GCF_000001405.40; Mus musculus, GCF_000001635.27; NCBI RefSeq assembly) using STAR (version 2.7.9a) on an Ubuntu 18.04 workstation. featureCounts function of the Rsubread package (version 2.0.3) was used for read counts count matrices. Downstream processing and analysis were performed in R (version 4.2.0) using the packages RNASeqQC, tidyverse, DESeq2 and edgeR. Quality control (QC) thresholds required ≥10 million aligned reads, 50–70% uniquely mapped reads, ≤10% rRNA reads, and ≥8000 detected genes per sample. In total, 37 libraries (human, n = 19; mouse, n = 18) passed QC (Fig. S2). Differential gene expression (DGE) analysis was performed using DESeq2. Genes with an absolute $\log_2$ fold change ≥ 1 and adjusted *P*-value ≤ 0.1 (Benjamini-Hochberg correction) were considered differentially expressed. Functional enrichment analysis was performed using clusterProfiler, enrichplot, R.utils, and DOSE, incorporating Kyoto Encyclopedia of Genes and Genomes (KEGG) pathways and Gene Set Enrichment Analysis (GSEA). Orthologous gene comparisons between species were conducted using the standalone Orthologous Matrix (OMA) tool with reference genomes from the OMA database. Scripts and detailed documentation for ortholog analysis are available on GitHub (https://github.com/Corleis-lab/Stei_et_al/tree/main/scripts). FASTQ files have been deposited in NCBI GEO under accession number GSE307006.

**Re-analysis of public scRNAseq and bulkRNAseq datasets**

For comparative analysis, previously published scRNAseq and bulkRNAseq datasets were re-analyzed for the expression of *NOS2, IFNγ* and *CXCL10* (Table S1). Data sets were downloaded according to the instructions provided in the respective publications, including accompanying metadata and cell-type annotations if available. Re-analysis was performed using R (version 4.3.2 or 4.3.3) and Seurat (version 5.3.0). Where cell-type annotations were not provided or required re-analysis, cell clusters were annotated using protocols described by Hou and Ji (2024) (2), assisted by code optimization through GPT Plus v4.0. All code used for dataset re-analysis is available on GitHub (https://github.com/Corleis-lab/Stei_et_al/tree/main/scripts).

**Statistical Analysis**

All statistical analyses were performed using GraphPad Prism (version 10.2.1), except for bulk RNA sequencing data, which were analyzed using DESeq2 (Wald-test and FDR correction; $\log_2$ fold change $\geq 1$ and adjusted p-value $\leq 0.1$) for differential expression and visualized using the ggplot2 R package. Due to limited sample sizes and sample types, we did not assume normal data distribution and therefore used non-parametric statistical tests throughout. Data are presented as medians with interquartile ranges and include individual data points for transparency unless stated otherwise in figure legends. For comparisons between two groups, we used two-sided Wilcoxon rank-sum tests (Mann-Whitney U test) or Wilcoxon signed-rank test for paired data sets. For comparisons involving more than two groups, Kruskal-Wallis (for independent groups) or Friedman test (for repeated measures) was applied, followed by Dunn's or Šídák's post hoc correction for multiple comparisons. Statistical significance was defined as $P < 0.05$ after adjustment for multiple hypothesis testing. For bulk RNA-seq analysis, differential gene expression was determined using DESeq2 with false discovery rate (FDR) adjustment using the Benjamini-Hochberg method. Gene set enrichment analysis (GSEA) was performed with FDR correction for pathway-level comparisons. Details on the number of biological replicates and the type of statistical test used are provided in each figure legend.

**Data and materials availability**

All raw and processed RNA sequencing data generated in this study have been deposited in NCBI Gene Expression Omnibus (GEO) under accession number GSE307006. Custom code for differential gene expression analysis, gene set enrichment, and ortholog mapping is available on GitHub https://github.com/Corleis-lab/Stei_et_al/tree/main/scripts.

All data are available in the main text or the supplementary materials. Additional raw data supporting the findings of this study, including ELISA, CFU, qRT-PCR, and Griess assay results, are available upon reasonable request from the corresponding author. Mycobacterium tuberculosis H37Rv::mCherry was generously provided by Stefan H.E. Kaufmann under a materials transfer agreement (MTA).

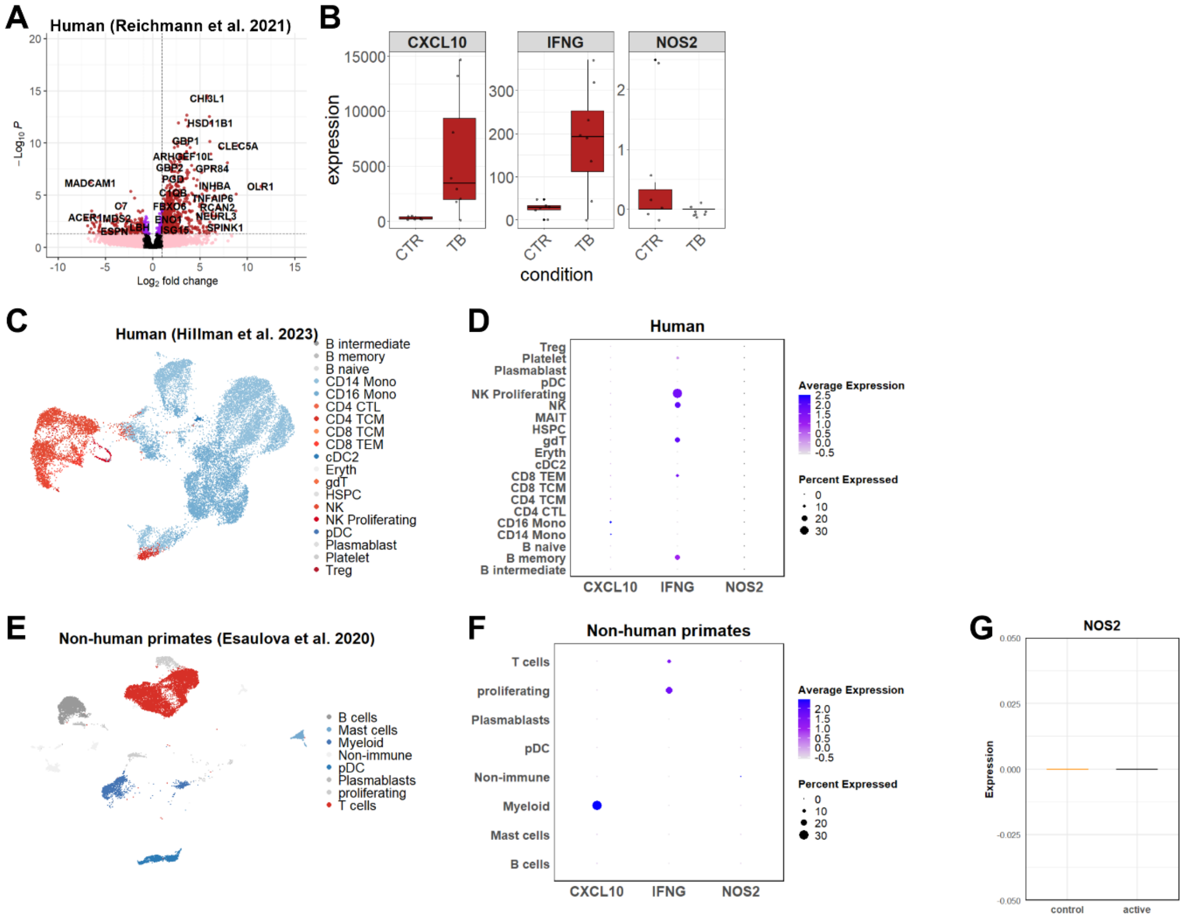

**Fig. S1.** ***NOS2*** **expression is absent in human and non-human primate myeloid cells despite IFNγ signaling.** (A) Bulk RNA-seq analysis from human lymph nodes (Reichmann et al., 2021; GSE174443) comparing TB patient samples and healthy controls shows elevated *IFNG* and *CXCL10* but minimal *NOS2* expression. (B) Normalized gene expression for *IFNG*, *CXCL10*, and *NOS2* across sample groups. (C) scRNA-seq from PBMCs (Hillmann et al., 2023; GSE214237) from TB patients reveals *CXCL10*+ monocyte clusters lacking *NOS2* expression. (D) Dot plot showing expression of *CXCL10*, *IFNG* and *NOS2* across major PBMC clusters. (E) UMAP of NHP granuloma cells from non-human primates (Esaulova et al., 2020; GSE149758) showing myeloid and lymphoid clusters. (F) Dot plots for *CXCL10*, *IFNG*, and *NOS2* expression across major NHP cell clusters. (G) Expression of *NOS2* across cell types in NHP granulomas (control samples versus active TB). Data confirm that *NOS2* is not expressed by myeloid cells despite active *IFNG* responses in human and NHP granulomas.

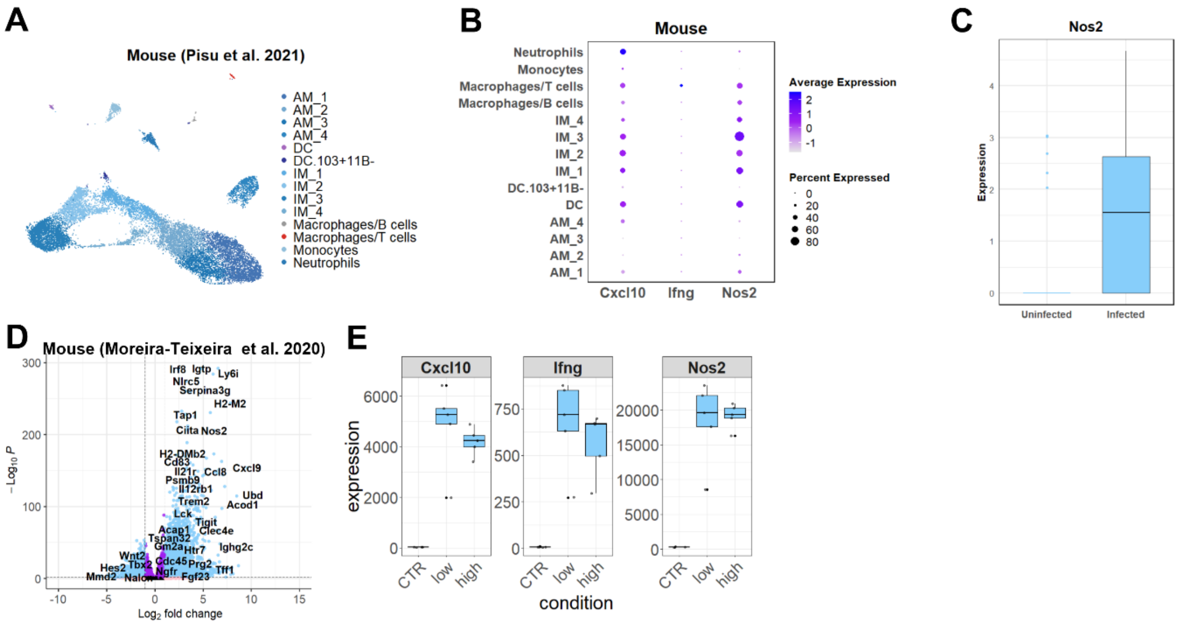

**Fig. S2. Species-specific *NOS2* expression in lung myeloid cells during *Mtb* or viral infection.** (A–C) Reanalysis of scRNA-seq data from mouse lungs infected with *Mtb* (Pisu et al., 2021; GSE167232). (A) UMAP showing major lung immune populations. (B) Dot plot of *Nos2*, *Cxcl10*, and *Ifng* expression across cell clusters; (C) Box plot showing *Nos2* expression in myeloid cells (uninfected versus infected). (D–E) Bulk RNA-seq data from *Mtb*-infected mice (Moreira-Teixeira et al., 2020; GSE137093). (D) Volcano plot of differentially expressed genes comparing control vs. high-dose infection, highlighting *Nos2* among strongly upregulated genes. (E) Box plots of *Cxcl10*, *Ifng*, and *Nos2* expression across control, low-dose, and high-dose groups.

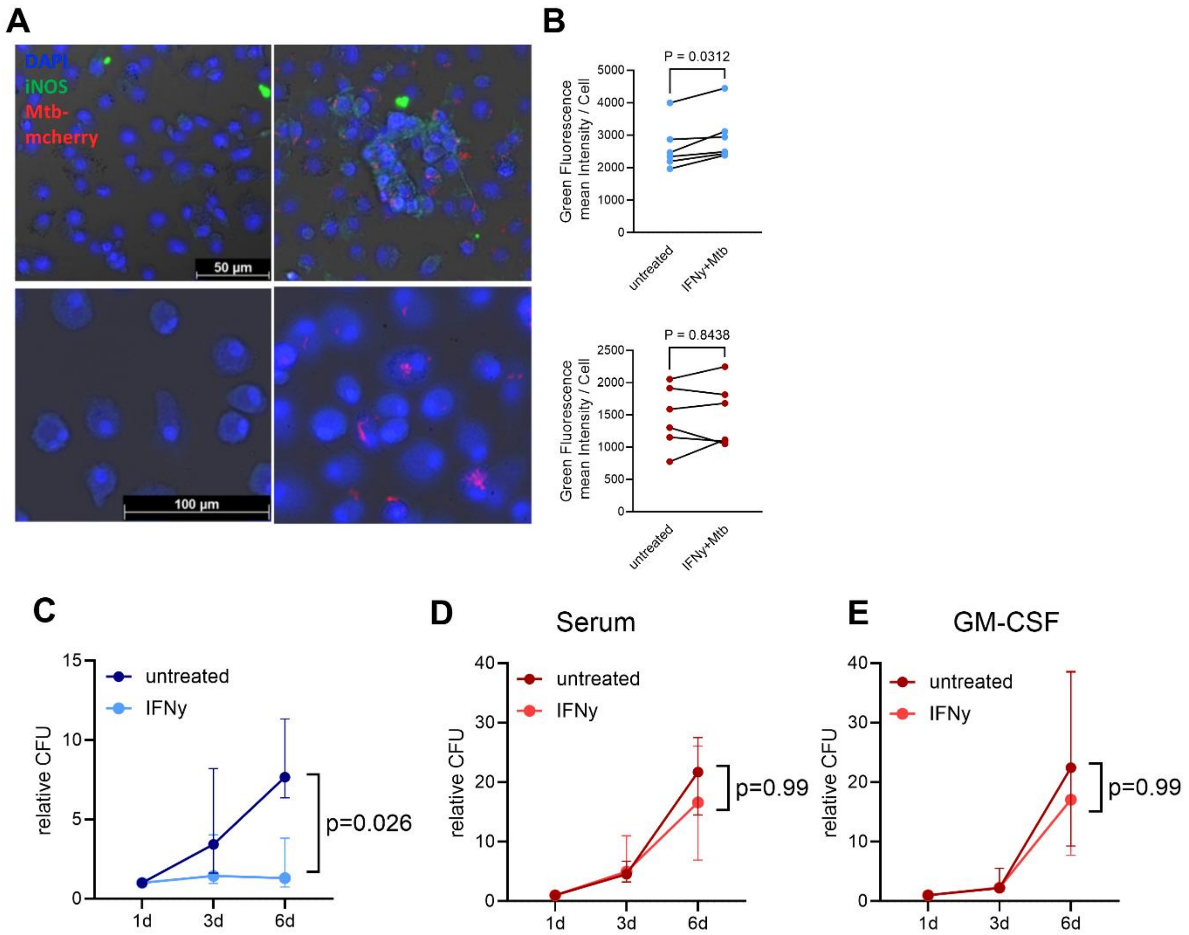

**Fig.S3: IFNγ activates human macrophages but does not reduce *Mtb* intracellular replication.** Human monocyte-derived macrophages were differentiated in either human serum or with recombinant GM-CSF, and mouse macrophages were derived from bone marrow in L929 cell-conditioned medium. (A) Representative immunofluorescence images of mouse (top) and human (bottom) macrophages infected with Mtb::mCherry (MOI 1-2) and stimulated with IFNγ (right) or left untreated / uninfected (left) for 24h (post infection). iNOS (green), nuclei (DAPI, blue), and Mtb (red) are shown. Scale bar = 50-100 µm. (B) Quantification of mean green fluorescence intensity (iNOS) per cell in mouse (blue) and human (red) macrophages; P-values determined by paired two-sided Wilcoxon test. (C–E) Mouse (blue) macrophages (n=6) and human (red) macrophages (serum-differentiated, n = 7; GM-CSF-differentiated, n = 3) were infected with *Mtb* H37Rv (MOI 0.1) following IFNγ stimulation. Intracellular bacterial burden was assessed by colony-forming unit (CFU) assay over time and shown relative to day 1 post-infection. *P*-values were determined by two-sided nonparametric tests. RM 2-way ANOVA with the Geisser-Greenhouse correction and post hoc Šídák's multiple comparisons test, showing median with interquartile range.

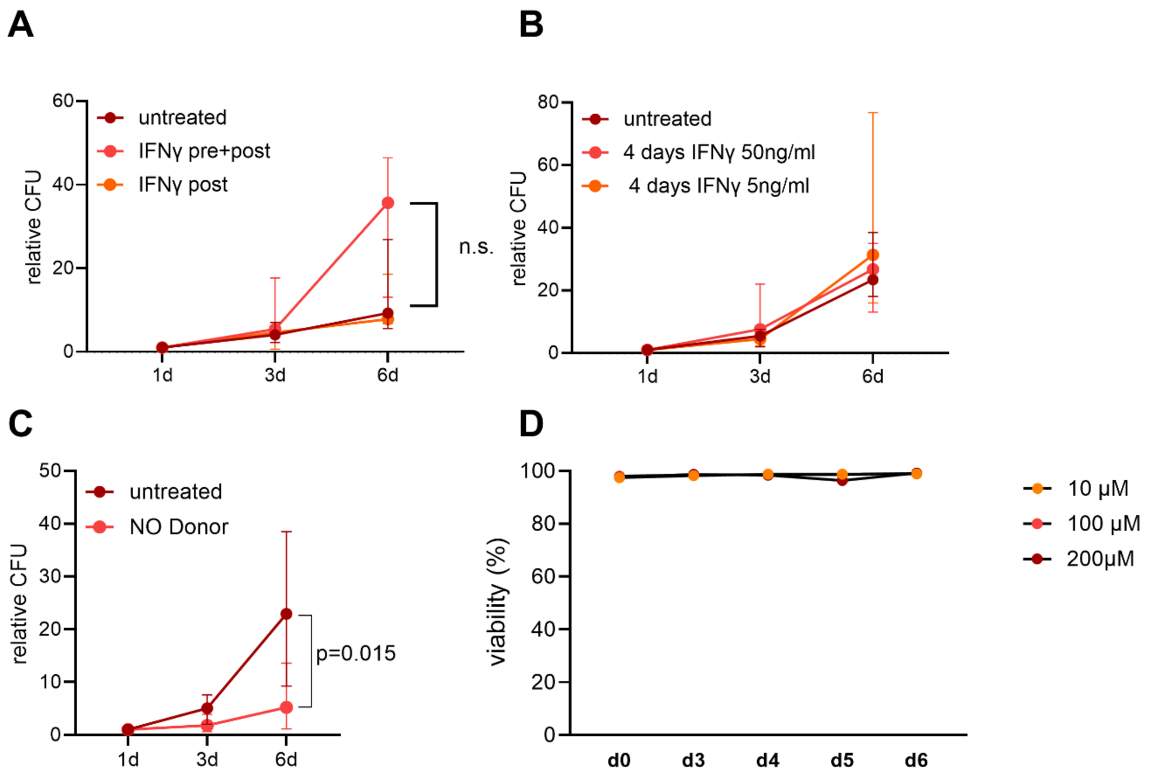

**Fig. S4. iNOS expression and nitric oxide production in IFNγ+*Mtb*-stimulated macrophages.**

(A) Relative CFU in macrophages treated with IFNγ either pre- and post-infection or post-infection only (n = 3). (B) Relative CFU in macrophages pre-treated with IFNγ for 4 days at two concentrations (5 ng/ml and 50 ng/ml) prior to infection (n = 5). (C) Relative CFU quantification of human macrophages infected with *Mtb* and treated with a NO donor (DETA NONOate) or left untreated, showing reduced bacterial burden at day 6 (n = 4); Statistical comparisons for CFU data were performed using repeated-measures two-way ANOVA with Geisser-Greenhouse correction and Šídák's post hoc test. Data shown as median ± range. (D) Viability of human macrophages (n=1) exposed to different concentrations (10–200 µM) of NO donor over 6 days.

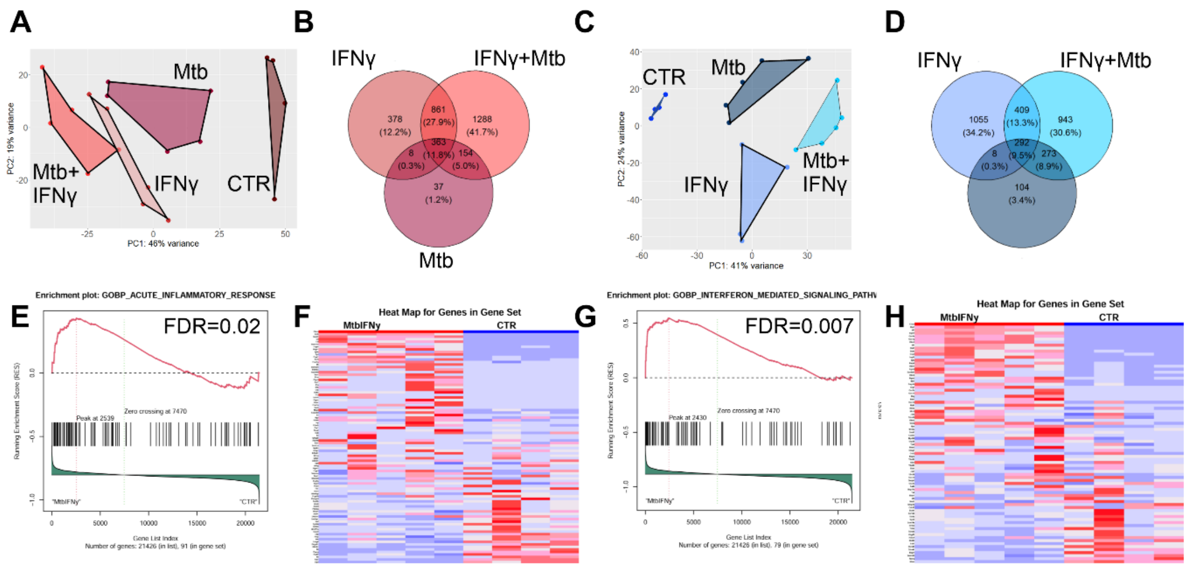

**Fig.S5. Bulk RNA sequencing reveals similar pro-inflammatory and IFNγ induced activation in human and mouse macrophages.** (A, B) Human (red) macrophages (differentiated in human serum) and (C-H) mouse (blue) bone marrow–derived macrophages were stimulated with IFNγ for 24 h, followed by *Mtb* infection (MOI 1–2) for 24 h. Controls were untreated or treated with IFNγ or *Mtb* alone. RNA was extracted and sequenced (Smart-seq2). (A, C) Principal component analysis (PCA) of transcriptomes from human (red; n = 19; CTR n = 4, IFNγ n = 5, *Mtb* n = 5 and *Mtb*+IFNγ n = 5) and mouse (blue; n = 18; CTR n = 4, IFNγ n = 4, *Mtb* n = 5 and *Mtb*+IFNγ n = 5) macrophages by condition. (B, D) Venn diagrams show the number, percentages and overlap of differentially expressed genes (DEGs) in each treatment group versus control. (E–H) Gene Set Enrichment Analysis (GSEA) for inflammatory and Interferon γ pathway enriched genes of DEGs from IFNγ+*Mtb* vs. control in mouse samples, including enrichment plots (E, G) and corresponding heatmaps (F, H). False discovery rates (FDRs) are indicated in each panel. Statistical analysis was performed using DESeq2 and clusterProfiler.

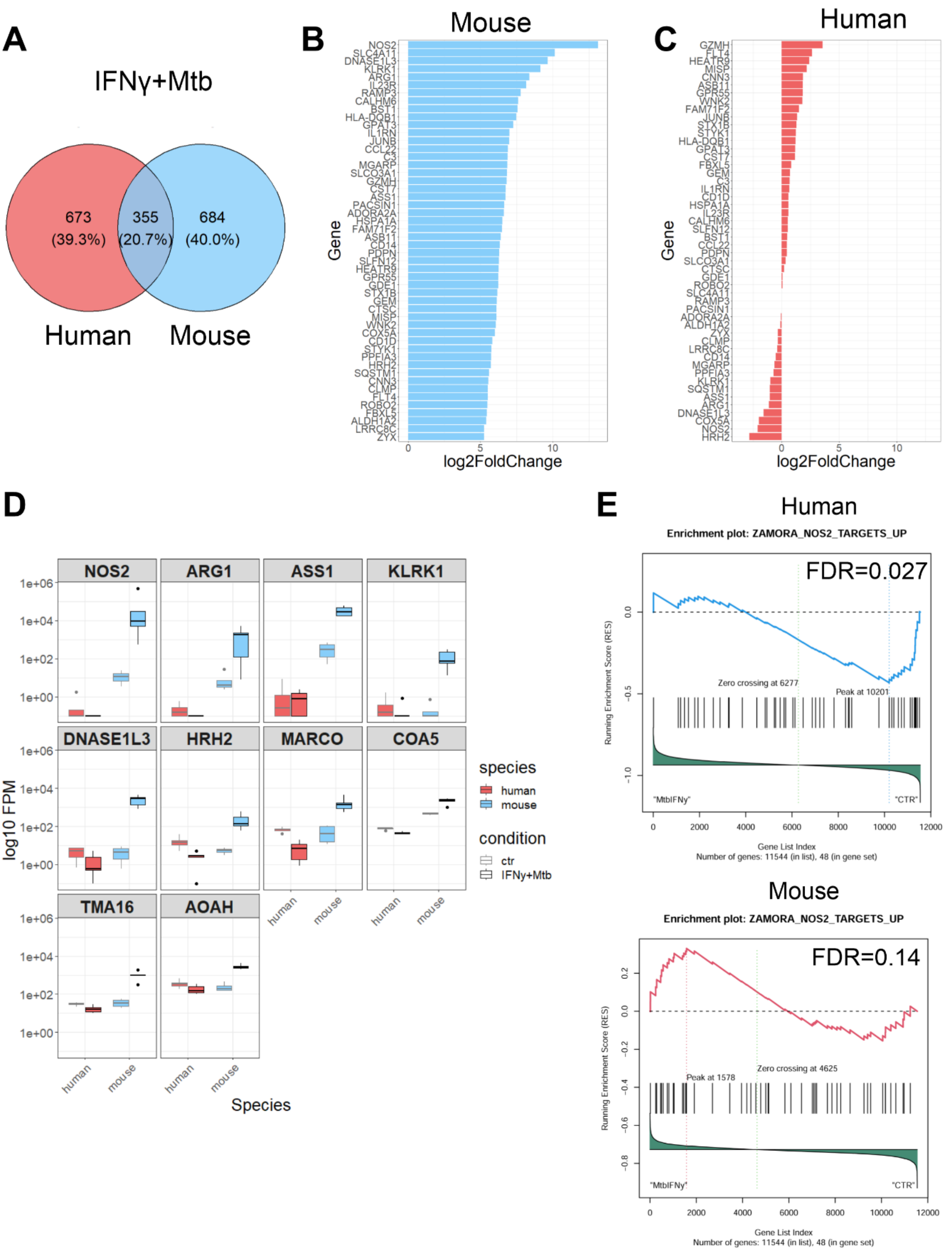

**Fig. S6. Interspecies ortholog comparison identifies *NOS2* as a key divergent gene between human and mouse macrophages.** (A) Venn diagram showing overlap of DEGs among 1712 one-to-one orthologous genes between IFNγ+*Mtb*-treated human (n = 5) and mouse (n = 5) macrophages versus untreated controls. (B–C) Top 50 DEGs (ranked by log2 fold change) in mouse (B) and corresponding log2 fold change in human macrophages (C). (D) Expression (log10 FPM) of selected ortholog genes with discordant regulation across species (box plots; human = red and mouse = blue). (E) GSEA using a reference gene set of *NOS2*-dependent genes in *NOS2*-/- hepatocytes transfected with human *NOS2 in vivo*. Adjusted *P*-values from GSEA are reported as FDR for enrichment in human (top) and mouse (bottom) macrophages.

A

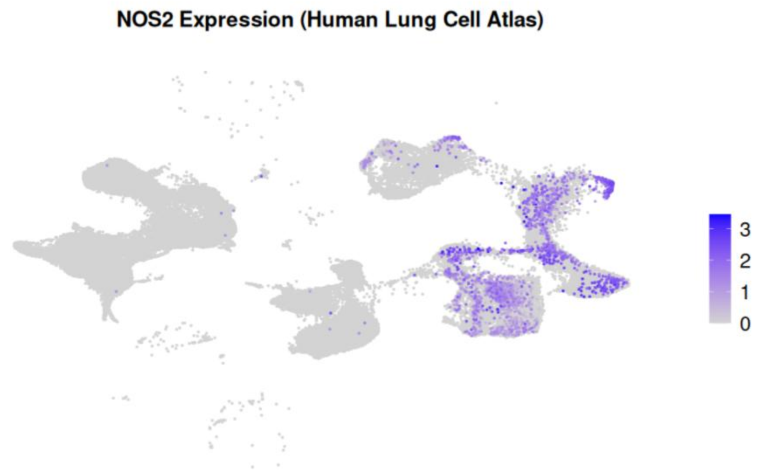

NOS2 Expression (Human Lung Cell Atlas)
3
2
1
0

B

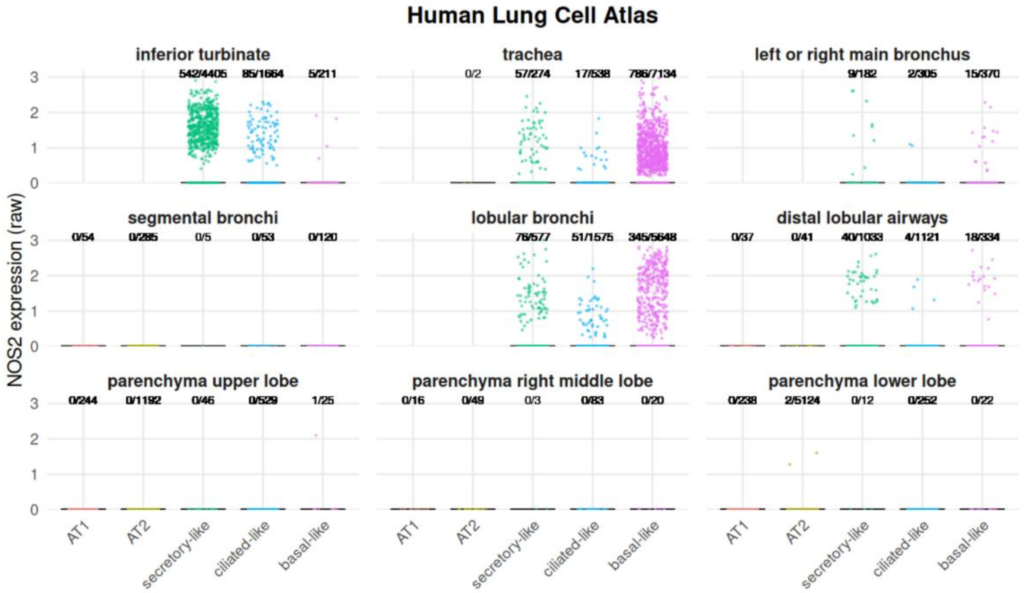

Human Lung Cell Atlas
NOS2 expression (raw)
inferior turbinate
542/4405 85/1664 5/211
trachea
0/2 57/274 17/538 786/7134
left or right main bronchus
9/182 2/305 15/370
segmental bronchi
0/54 0/285 0/5 0/53 0/120
lobular bronchi
76/577 51/1575 345/5648
distal lobular airways
0/37 0/41 40/1033 4/1121 18/334
parenchyma upper lobe
0/244 0/1192 0/46 0/529 1/25
parenchyma right middle lobe
0/16 0/49 0/3 0/83 0/20
parenchyma lower lobe
0/238 2/5124 0/12 0/252 0/22
AT1
AT2
secretory-like
ciliated-like
basal-like

C

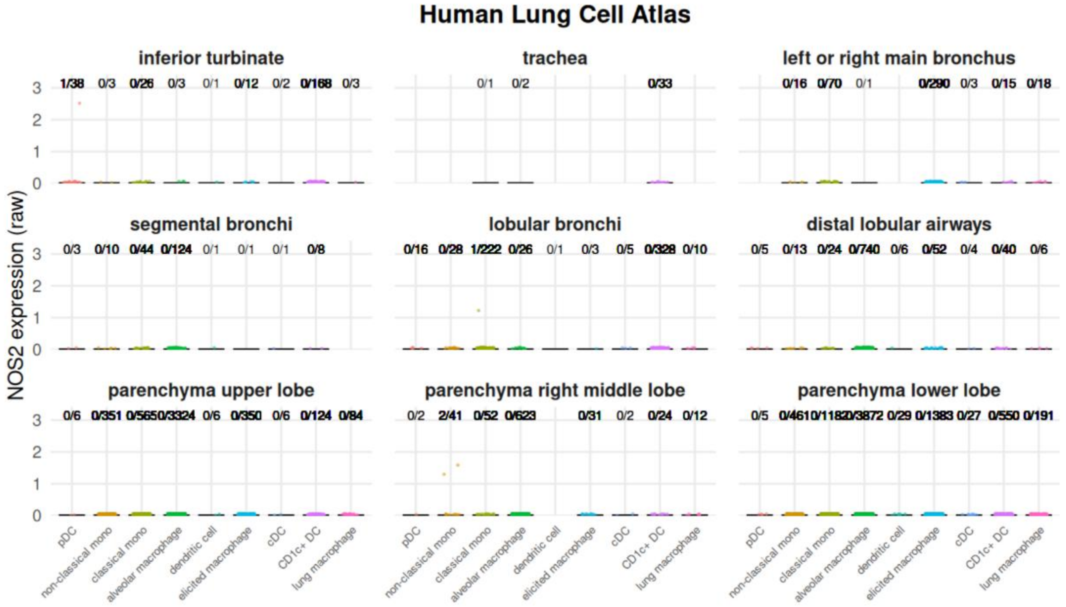

Human Lung Cell Atlas
NOS2 expression (raw)
inferior turbinate
1/38 0/3 0/26 0/3 0/1 0/12 0/2 0/168 0/3
trachea
0/1 0/2 0/33
left or right main bronchus
0/16 0/70 0/1 0/290 0/3 0/15 0/18
segmental bronchi
0/3 0/10 0/44 0/124 0/1 0/1 0/1 0/8
lobular bronchi
0/16 0/28 1/222 0/26 0/1 0/3 0/5 0/328 0/10
distal lobular airways
0/5 0/13 0/24 0/740 0/6 0/52 0/4 0/40 0/6
parenchyma upper lobe
0/6 0/351 0/5650/3324 0/6 0/350 0/6 0/124 0/84
parenchyma right middle lobe
0/2 2/41 0/52 0/623 0/31 0/2 0/24 0/12
parenchyma lower lobe
0/5 0/4610/1182 0/3872 0/29 0/1383 0/27 0/550 0/191
pDC
non-classical mono
classical mono
alveolar macrophage
dendritic cell
elicited macrophage
cDC
CD1c+ DC
lung macrophage

**Fig. S7. *NOS2* expression is predominantly detected in secretory epithelial cells of the upper airways in human lung tissue.** (A) UMAP projection of single-cell RNA-seq data from the Human Lung Cell Atlas (Sikkema *et al*., 2023; https://data.humancellatlas.org/hca-bio-networks/lung/atlases/lung-v1-0), colored by NOS2 expression levels. (B–C) *NOS2* expression by cell type and anatomical lung region. (B) *NOS2* expression (raw counts) across epithelial cell subsets (AT1, AT2, secretory-like, ciliated-like, basal-like) in different airway and parenchymal compartments. (C) *NOS2* expression across myeloid cell populations (e.g., non-classical monocytes, cDCs, pDCs) from the same anatomical compartments. Ratios indicate number of *NOS2*$^{+}$ cells per total cells in each category.

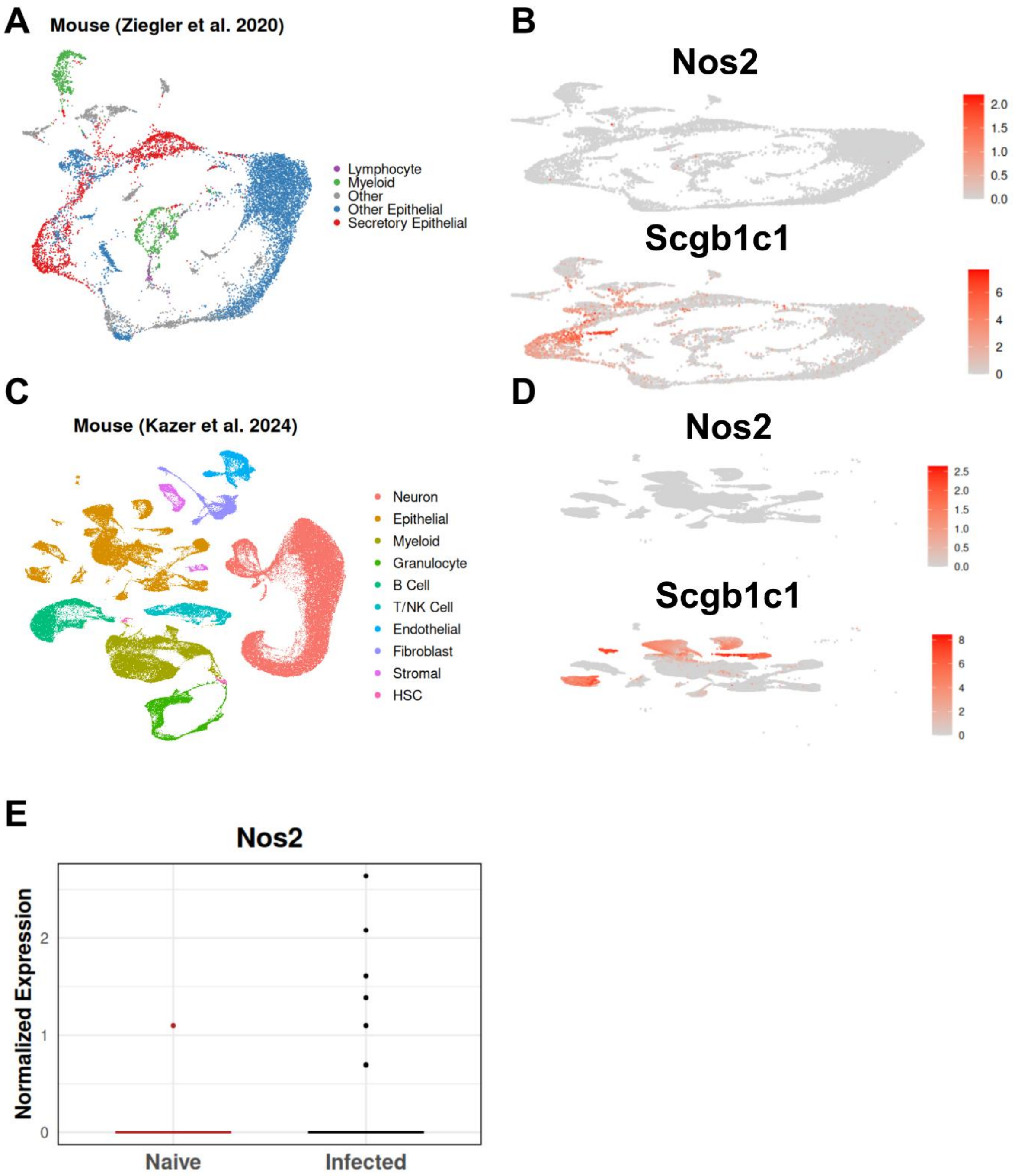

**Fig. S8. Mouse nose mucosa epithelial cells have low expression of *Nos2* in secretory epithelial cells.** (A) UMAP projection of scRNA-seq data from the mouse nasal epithelium (Ziegler et al., 2020; SCP832), annotated by cell type. (B) Feature plots showing *Nos2* and *Scgb1c1* expression across cell types in (A). (C) UMAP of an independent mouse lung dataset (Kazer et al., 2024; SCP2216), annotated by major cell populations. (D) Feature plots of *Nos2* and *Scgb1c1* expression in the dataset shown in (C). (E) Box plot of normalized Nos2 expression in secretory epithelial cells from naïve and H1N1-infected animals.

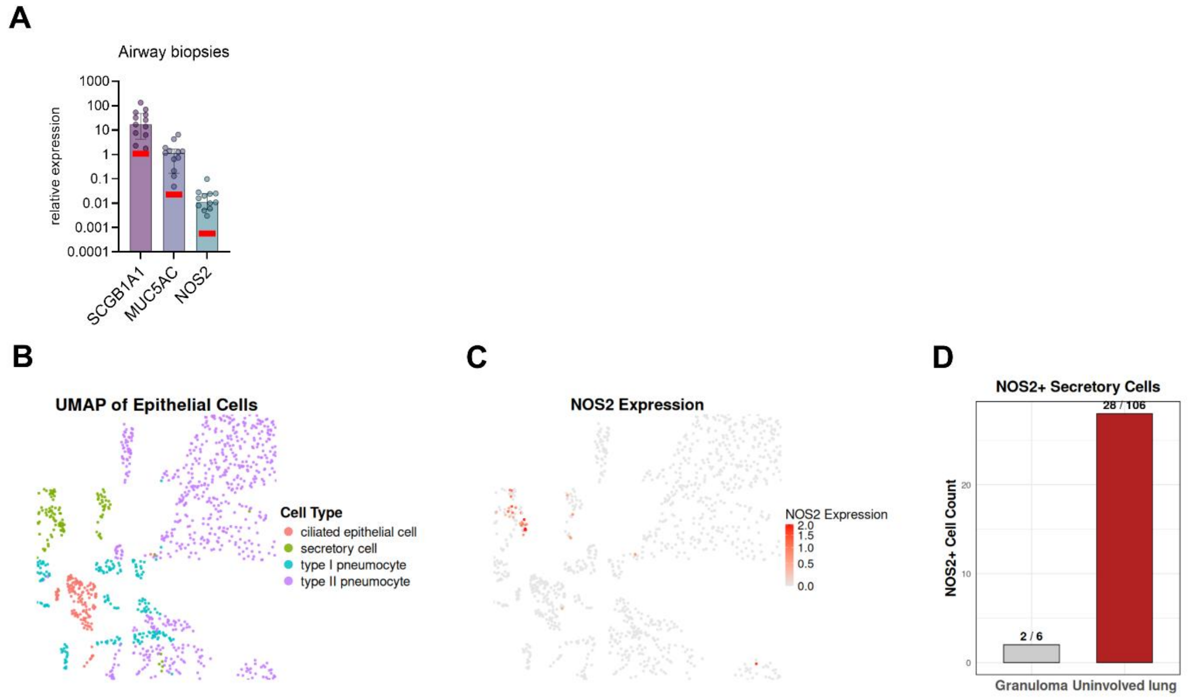

**Fig. S9.** ***NOS2* is expressed in secretory epithelial cells from uninvolved lung tissue in non-human primates with M. tuberculosis infection.** (A) relative RNA expression (2^- ΔCt) of *SCG1A1*, *MUC5AC* and *NOS2* in human airway biopsy samples (n=12). Data are presented as median with interquartile range and individual data points. Red bar indicates background expression in two BAL sample. (B) UMAP of epithelial cells from macaque granuloma and uninvolved lung tissue, colored by cell type (ciliated epithelial cell, secretory cell, type I pneumocyte, type II pneumocyte), adapted from SCP806, Broad Institute Single Cell Portal. (C) Feature plot showing *NOS2* expression across epithelial cells. (D) Quantification of NOS2$^+$ secretory cells by tissue origin. Numbers above bars indicate *NOS2*$^+$/total secretory cells.

**Table S1.** Overview public single cell and bulk RNAseq data sets used for re-analysis

| Species | First author | Data link | Data type | Pathogen | Sample type |
|---|---|---|---|---|---|
| Human | Wang *et al.* | https://www.ncbi.nlm.nih.gov/geo/query/acc.cgi?acc=GSE192483 | scRNAseq | *Mtb* | lung resections from 4 TB patients |
| Human | Reichmann *et al.* | https://www.ncbi.nlm.nih.gov/geo/query/acc.cgi?acc=GSE174443 | bulkRNAseq | *Mtb* | mediastinal and neck lymph node granuloma resections |
| Human | Hillmann et al. | https://www.ncbi.nlm.nih.gov/geo/query/acc.cgi?acc=GSE214237 | scRNAseq | *Mtb* | PBMC samples of individuals with active tuberculosis |
| Non-human primates | Gideon *et al.* | https://www.ncbi.nlm.nih.gov/geo/query/acc.cgi?acc=GSE200151 | scRNAseq | *Mtb* | lung granuloma 4 & 10 weeks after Mtb infection |
| Non-human primates | Esaulova *et al.* | https://www.ncbi.nlm.nih.gov/geo/query/acc.cgi?acc=GSE149758 | scNRAseq | *Mtb* | CD45+ enriched samples from macaca mulatta lungs |
| Mouse | Kang *et al.* | https://www.ncbi.nlm.nih.gov/geo/query/acc.cgi?acc=GSE167650 | scRNAseq | *Mtb* | CD45+ immune cells from infected mouse lungs |
| Mouse | Pisu *et al.* | https://www.ncbi.nlm.nih.gov/geo/query/acc.cgi?acc=GSE167232 | scRNAseq | *Mtb* | mouse lungs at 3 weeks post infection |
| Mouse | Moreira-Teixeira *et al.* | https://www.ncbi.nlm.nih.gov/geo/query/acc.cgi?acc=GSE137093 | bulkRNAseq | *Mtb* | whole lung samples |
| Human | Liao *et al.* | https://www.ncbi.nlm.nih.gov/geo/query/acc.cgi?acc=GSE145926 | scRNAseq | SARS-CoV-2 | BAL fluid from patients with mild or severe COVID-19 |
| Human | Human Lung Cell Atlas | https://data.humancellatlas.org/hca-bio-networks/lung/atlases/lung-v1-0 | scRNAseq | N/A | 4 respiratory control tissues from patients with various diseases |

| | | | | | |
|---|---|---|---|---|---|
| Human | Ziegler *et al.* | https://singlecell.broadinstitute.org/single_cell/study/SCP1289/impaired-local-intrinsic-immunity-to-sars-cov-2-infection-in-severe-covid-19#study-summary | scRNAseq | SARS-CoV-2 | Nasopharyngeal swabs |
| Mouse | Ziegler *et al.* | https://singlecell.broadinstitute.org/single_cell/study/SCP832/murine-nasal-mucosa-after-intranasal-interferon-exposure#study-summary | scRNAseq | N/A | Nasal mucosa 12 h after IFNα treatment |
| Mouse | Kazer *et al.* | https://singlecell.broadinstitute.org/single_cell/study/SCP2216/primary-nasal-viral-infection-rewires-the-tissue-scale-memory-response-primary-infection-dataset#study-summary | scRNAseq | H1N1 | murine nasal mucosa |
| Non-human primates | Ziegler *et al.* | https://singlecell.broadinstitute.org/single_cell/study/SCP806/epithelial-cells-in-nhp-mtb-granuloma-and-uninvolved-lung | scRNAseq | *Mtb* | Epithelial cell enriched lung samples from infected animals |

**Table S2.** Summary and sequence of all primer pairs

| Species | Gene Symbol | Gene ID | Primer sequence | Amplicon size / bp |
|---|---|---|---|---|
| **Human** | *CXCL10* | 3627 | F GTGGCATTCAAGGAGTACCTC | 198 |
| | | | R TGATGGCCTTCGATTCTGGATT | |
| | *GAPDH* | 2597 | F CCTCAACGACCACTTTGTCA | 103 |
| | | | R TTACTCCTTGGAGGCCATGT | |
| | *NOS2* | 4843 | F TGCTCAGCTCATCCGCTATG | 78 |
| | | | R GTGAATTCCACGTTGGCAGG | |
| | *NOS2* | 4843 | F CCACTGCCCGGGAAATGTTTG | 70 |
| | | | R CTGATGTTGCCATTGTTGGTGG | |
| | *RPS13* | 6207 | F CGAAAGCATCTTGAGAGGAACA | 87 |
| | | | R TCGAGCCAAACGGTGAATC | |
| | *SCGB1A1* | 7356 | F CAAAAGCCCAGAGAAAGCATC | 93 |
| | | | R CAGTTGGGGATCTTCAGCTTC | |
| | *MUC5AC* | 4586 | F GCCTTCACTGTACTGGCTGAG | 149 |
| | | | R TGGGTGTAGATCTGGTTCAGG | |
| **Mouse** | *Cxcl10* | 15945 | F CCAAGTGCTGCCGTCATTTTC | 157 |
| | | | R GGCTCGCAGGGATGATTTCAA | |
| | *Gapdh* | 14433 | F CTACCCCCAATGTGTCCGTC | 151 |
| | | | R AAGTCGCAGGAGACAACCTG | |
| | *Nos2* | 18126 | F CTTGGTGAAGGGACTGAGCTG | 116 |
| | | | R TCCAAATCCAACGTTCTCCGT | |

**Table S3.** qPCR cycler program

| Step | Temperature in °C | Time | Cycles |
|---|---|---|---|
| Warm Up | *25* | *2 min* | 1 |
| | *95* | *2 min* | |
| PCR | *95* | *15 sec* | 40 |
| | *60* | *30 sec* | |
| | Plate read | | |
| Melt curve | *95* | *15 sec* | 1 |
| | *60* | *1 min* | |
| | 60 → 95 (0,15 °C/s) | / | |
| | + Plate read | | |
| | *95* | *15 sec* | |

**Table S4.** Overview of all master mixes and PCR programs for SmartSeq 2 libraries

| Mastermix #1 (green label) | |
|---|---|
| **Reagent** | **for 1 sample (µl)** |
| 3' RT primer (100 uM) | 0,1 |
| dNTP Mix (10 mM each NTP) | 1 |
| Rnase inhibitor (40 U/uL) | 0,1 |
| H2O | 2,8 |
| **Total** | **4** |

| Mastermix #2 (red label) | |
|---|---|
| **Reagent** | **for 1 sample (µl)** |
| 5X Maxima RT buffer | 2 |
| Betaine (5M) | 2 |
| MgCl2 (1M) | 0,09 |
| TSO (100 uM) | 0,1 |
| Maxima Rnase H-minus RT (200 U/uL) | 0,1 |
| Rnase inhibitor (40 U/uL) | 0,25 |
| H2O | 2,46 |
| **Total** | **7** |

| Mastermix #3 (white label) | |
|---|---|
| **Reagent** | **for 1 sample (µl)** |
| IS PCR Primer (100 uM) | 0,05 |
| 2X KAPA HiFi HotStart Ready Mix | 12,5 |
| H2O | 1,45 |
| **Total** | **14** |

| Nextera reagents | |
|---|---|
| **Reagent** | **for 1 sample (µl)** |
| TD buffer (Tagement DNA buffer) | 2,5 |
| ATM enzyme (Amplicon Tagment Mix) | 1,25 |
| NT buffer | 1,25 |
| Nextera Indexing Plate | 2,5 |
| Nextera PCR Mastermix (NPM) | 3,75 |

| PCR Protocol #1 | | | |
|---|---|---|---|
| **Info** | **Cycles** | **Temperature [°C]** | **Time [min]** |
| Lid temp | - | 105 | - |
| | - | 42 | 90 |
| PCR | 10 | 50 | 2 |

| | | | |
|---|---|---|---|
| | | 42 | 2 |
| Heat Inactivation | - | 70 | 15 |
| Hold | - | 4 | Infinity |
| **PCR Protocol #2** | | | |
| **Info** | **Cycles** | **Temperature [°C]** | **Time [min]** |
| Lid temp | - | 105 | - |
| cDNA denaturation | - | 98 | 3 |
| | | 98 | 15s |
| | | 67 | 20s |
| PCR | 21 | 72 | 6 |
| Final extention | - | 72 | 5 |
| Hold | - | 4 | Infinity |
| **PCR Protocol #3** | | | |
| **Info** | **Cycles** | **Temperature [°C]** | **Time [min]** |
| Lid temp | - | 105 | - |
| Tn5 cutting | - | 55 | 10 |
| Hold | - | 4 | Infinity |
| **PCR Protocol #4** | | | |
| **Info** | **Cycles** | **Temperature [°C]** | **Time [min]** |
| Lid temp | - | 105 | - |
| Elongation completion | - | 72 | 3 |
| cDNA denaturation | - | 95 | 30s |
| | | 95 | 10s |
| | | 55 | 30s |
| PCR | 12 | 72 | 1 |
| Final extention | - | 72 | 5 |
| Hold | - | 4 | Infinity |
